# Cleavage site-directed antibodies reveal the prion protein in humans is shed by ADAM10 at Y226 and associates with misfolded protein deposits in neurodegenerative diseases

**DOI:** 10.1101/2023.11.30.569390

**Authors:** Feizhi Song, Valerija Kovac, Behnam Mohammadi, Lisa Littau, Franka Scharfenberg, Andreu Matamoros Angles, Ilaria Vanni, Mohsin Shafiq, Leonor Orge, Giovanna Galliciotti, Salma Djakkani, Luise Linsenmeier, Maja Černilec, Katrina Hartman, Sebastian Jung, Jörg Tatzelt, Julia E. Neumann, Markus Damme, Sarah K. Tschirner, Stefan F. Lichtenthaler, Matthias Schmitz, Inga Zerr, Berta Puig, Eva Tolosa, Isidro Ferrer, Tim Magnus, Marjan S. Rupnik, Diego Sepulveda-Falla, Jakob Matschke, Lojze M. Šmid, Mara Bresjanac, Olivier Andreoletti, Susanne Krasemann, Simote T. Foliaki, Romolo Nonno, Christoph Becker-Pauly, Cecile Monzo, Carole Crozet, Cathryn L. Haigh, Markus Glatzel, Vladka Curin Serbec, Hermann C. Altmeppen

## Abstract

Proteolytic cell surface release (‘shedding’) of the prion protein (PrP), a broadly expressed GPI-anchored glycoprotein, by the metalloprotease ADAM10 impacts on neurodegenerative and other diseases in animal and *in vitro* models. Recent studies employing the latter also suggest shed PrP (sPrP) to be a ligand in intercellular communication and critically involved in PrP-associated physiological tasks. Although expectedly an evolutionary conserved event, and while soluble forms of PrP are present in human tissues and body fluids, neither proteolytic PrP shedding and its cleavage site nor involvement of ADAM10 or the biological relevance of this process have been demonstrated for the human body thus far. In this study, cleavage site prediction and generation (plus detailed characterization) of sPrP-specific antibodies enabled us to identify PrP cleaved at tyrosin 226 as the physiological and strictly ADAM10-dependent shed form in humans. Using cell lines, neural stem cells and brain organoids, we show that shedding of human PrP can be stimulated by PrP-binding ligands without targeting the protease, which may open novel therapeutic perspectives. Site-specific antibodies directed against human sPrP also detect the shed form in brains of cattle, sheep and deer, hence in all most relevant species naturally affected by fatal and transmissible prion diseases. In human and animal prion diseases, but also in patients with Alzheimer’s disease, sPrP relocalizes from a physiological diffuse tissue pattern to intimately associate with extracellular aggregates of misfolded proteins characteristic for the respective pathological condition. Findings and research tools presented here will accelerate novel insight into the roles of PrP shedding (as a process) and sPrP (as a released factor) in neurodegeneration and beyond.

## Introduction

Proteolytic processing is of utmost importance for regulating certain proteins’ physiological functions, yet also plays important roles in diverse pathological conditions ^1^. For some “multifunctional” proteins, conserved cleavages by endogenous proteases do not just simply reflect a start of inactivation or catabolic degradation, but rather represent the impetus for functional diversity, regulation and effects mediated by the resulting fragments. The prion protein (PrP), a membrane-anchored glycoprotein with high (though not exclusive) expression in the nervous system, may be considered as a multifunctional protein, at least in view of the variety of suggested physiological tasks ^2–5^. In contrast, its key pathological role in fatal and transmissible neurodegenerative prion diseases such as Creutzfeldt-Jakob disease (CJD), where it serves as the critical substrate for a templated and progressive misfolding and aggregation process resulting in neuronal death and brain vacuolization, is firmly established ^6–9^. And a relevant role of PrP as a neuronal cell surface receptor for other toxic protein conformers in more common neurodegenerative diseases (such as Alzheimer’s (AD) or Parkinson’s disease) is being increasingly recognized ^10–16^.

Some conserved endogenous cleavage events within PrP have been identified over the years, yet their biological relevance is just starting to be understood as more systematic studies are being conducted ^5,17–20^. This also applies to the constitutive, membrane-proximate shedding by the metalloprotease ADAM10 ^21–23^, which is of particular interest as it releases nearly full-length PrP, i.e., shed PrP (sPrP), into the extracellular space while leaving only the GPI anchor and a few amino acids behind at the plasma membrane. This cleavage not only is a critical mechanistic part of a compensatory network ensuring cellular PrP homeostasis ^24^. It also impacts on neurodegenerative diseases by reducing cell surface PrP as a relevant receptor for (neuro)toxic protein assemblies ^25,26^. Moreover, once released into the extracellular space and interstitial fluid, sPrP may block, detoxify and sequester harmful oligomers into less toxic deposits ^26,27^, as observed earlier for recombinant PrP (recPrP) *in vitro* or transgenically expressed PrP dimers serving as a proxy for physiologically shed PrP ^28–33^. Fittingly, in prion disease mouse models, ADAM10 expression correlates with incubation and survival time ^34,35^; and sPrP levels inversely correlate with PrP^Sc^ formation ^26,27,35–37^. This collectively supports the notion that soluble PrP forms like sPrP may indeed act as “prion replication antagonists” ^28,38–40^. Recent studies in transgenic mice also highlight an influence of the ADAM10-mediated PrP shedding on prion strain characteristics and resulting aggregate morphology ^41,42^. Hence, manipulation of this particular proteolytic process appears to be a promising option against neurodegenerative diseases ^25–27^. We have recently uncovered a substrate-specific approach to this in murine cells and tissue using PrP-directed ligands ^27^, thereby avoiding general targeting of ADAM10 and, hence, likely side effects by affecting its manifold substrates in different organs ^43,44^.

Apart from neurodegenerative conditions, recent studies on potential biological roles of sPrP suggest it to act as a ligand inducing effects in different recipient cell types, regulating cellular differentiation, homeostasis, morphogenesis and immunological processes ^45–49^. Depending on the pathophysiological context, sPrP may also play detrimental roles as suggested by studies linking it with elevated inflammatory responses in HIV neuropathogenesis ^50^ as well as trophic effects or drug-resistance in cancer ^51,52^. In sum, these studies indicate sPrP as a novel factor in intercellular communication with likely cell-, tissue- and context-dependent consequences, and highlight the need for further investigations on protective as well as potentially harmful functions of sPrP in the nervous system and beyond. However, as those studies mainly used recPrP as a non-physiological proxy for bona fide sPrP, this may pose limitations regarding interpretation of the findings, as –for instance– differences in structure or glycosylation state may very well influence the experimental outcome ^26^.

Though providing critical initial insight, most studies on biological roles played by sPrP were limited to *in vitro* or animal models, thus leaving an important gap of knowledge for the human system. Moreover, when it comes to systematically investigating effects of PrP shedding and intrinsic functions of sPrP in meaningful models, there is at least one major hurdle: In brain or other tissue samples, and even in body fluids, reliable detection of sPrP using pan-PrP antibodies is difficult given the vastly exceeding and masking amounts of full-length membrane-bound PrP (either cell-associated or on extracellular vesicles (EVs)) with almost similar molecular weight and a current lack of discriminating antibodies ^19,26^. For the murine system, we have recently overcome this problem by generating cleavage site-specific antibodies for sPrP (not detecting its uncleaved GPI-anchored precursor ^24^) based on the previously published cleavage site ^22^. For the human body, however, strictly speaking neither the proteolytic C-terminal shedding in general, nor clear (and sole) involvement of ADAM10 in this process or the respective cleavage site within PrP have been convincingly shown or identified to date. This is surprising given that constitutive and reactive PrP release by different human cell types with likely (patho)physiological relevance, and presence of respective soluble, nearly full-length PrP forms in human brain tissue, cerebrospinal fluid (CSF) and blood have been shown manifold over the last three decades ^53–59^. Moreover, reports on the existence of nearly full-length, C-terminally truncated PrP variants in some prion disease patients, caused by stop mutations ^60^ or as yet unknown reasons ^61–65^, were followed by speculations on a potential link between such fragments and proteolytic shedding ^66^.

Supported by structural cleavage site predictions, we generated, characterized and compared poly- and monoclonal antibodies specifically detecting physiological C-terminally shed PrP in human samples. By demonstrating both the cleavage site at position Y226↓Q227 and functionality of these antibodies in different paradigms, we also show clear and exclusive ADAM10 dependency of respective signals. Further, as previously demonstrated in murine samples ^27^, we show that shedding can be stimulated by PrP-directed antibodies in different models of human origin and discuss its therapeutic feasibility. We assess PrP shedding in central nervous system (CNS) tissue, CSF and various cell types using different technical approaches. Using heterologous expression models, we also show that murine ADAM10 is able to cleave human PrP at the ‘human’ cleavage site, while human ADAM10 on murine PrP employs G227↓R228, the proper cleavage site in mice and rats. Strikingly, because of similar C-terminal sequences, the antibodies for human sPrP presented herein also detect shed PrP fragments in some of the most prion disease-relevant animal species such as cattle, sheep/goats, and deer, thus likely enabling future studies on the relevance of PrP shedding in bovine spongiform encephalopathy (BSE), Scrapie and chronic wasting disease (CWD), respectively. Lastly, we address PrP shedding in samples of patients affected by neurodegenerative diseases including CJD and AD, and demonstrate that sPrP, in the presence of proteopathic aggregates, is redistributed from an originally nondescript and diffuse to a plaque-associated pattern, thus warranting further studies on the proposed role of sPrP in blocking and sequestering extracellular toxic oligomers into potentially less harmful deposits. Moreover, we suggest future investigations to assess sPrP’s potential as an (easily) accessible biomarker in body fluids. In conclusion, we here provide novel information and research tools to study a formerly underestimated yet increasingly appreciated and evolutionary conserved proteolytic cleavage event on a key player in neurodegenerative proteinopathies, with therapeutic potential and biological relevance probably not being restricted to the CNS.

## Results

### Cleavage site prediction and targeted antibody production

Due to alterations in the C-terminal amino acid sequence between human and rodent PrP and lack of a glycine in a similarly membrane-proximate position as G227 (i.e. the P1 cleavage site and neo-C-terminus of sPrP in mice and rats ^22,67^), a different cleavage site for the shedding of human PrP was to be expected. Accordingly, our sPrP^G227^ antibody previously generated for rodent sPrP ^24^ is ineffective towards the human protein. We therefore combined educated guessing and cleavage site prediction based on available sequence and structural data for human PrP (^217^YERESQAYYQRGS^230^) as potential substrate (Source: www.uniprot.org; Major prion protein [*Homo sapiens*], ID: P04156) and human ADAM10’s catalytic domain ^68^.

Although modelling the C-terminal sequence of PrP within the catalytic domain of ADAM10 is difficult (e.g., because of uncertainty regarding structural constraints imposed by PrP’s GPI anchor), our modelling with PEP-FOLD3 ^69^ and FlexPepDock ^70^ suggested the PrP tyrosine at 226 (Y226) as a possible P1 cleavage site within ^217^YERESQAYY^226^↓QRGS^230^ (pink peptide), as shown by superposition with the enzyme-product complex of the C-terminus (^642^FMRCRLVDADGPLG^655^; yellow peptide) of adjacent ADAM10 subunits captured in the crystal structure of ADAM10 ^68^ (**Fig. 1a** (I and IV)). This conformation is possibly stabilized by R228 building a salt bridge with ADAM10’s E298 (**Fig. 1a** (II, III and V)) while ^217^YERESQAYY^226^ (the conceivable C-terminal ending of newly formed sPrP) is being released from the catalytic center (Fig. 1a (IV)). Albeit PrP may not be regarded as an ‘ideal’ substrate (note that the vast majority of ADAM substrates are transmembrane proteins) and alternative cleavage sites would have been conceivable from a structural perspective, certain residues were in agreement with published cleavage site preferences identified with substrate libraries for recombinant ADAM10 using *quantitative proteomics for the identification of cleavage sites* (Q-PICS; ^71^) or *terminal amine isotopic labelling of substrates* (TAILS; ^72^) (**Fig. 1b**). These data demonstrated that amino acid preferences of ADAM10 around the cleavage site slightly differ for peptide and protein substrates. Due to its GPI anchor and N-glycans, mature cellular PrP exhibits additional molecular properties that certainly impact ADAM10 cleavage. Hence, although not in line with all ‘most preferred’ amino acids identified in the previous studies, cleavage of PrP by ADAM10 at Y226↓Q227 fits to residues A224 (in P3), Y226 (P1) and G229 (P3’) (for PICS) as well as S222 (in P5) and G229 (P3’) (for TAILS), while no disfavored residues are present. Moreover, the distance of ∼20-25 Å between the potential cleavage site and the plasma membrane (here mostly determined by PrP’s GPI anchor ^73^) is in line with the membrane-proximity preferred by ADAM10 in complex with its regulator tetraspanin 15 ^74^ (which is involved in PrP shedding ^75^).

**Figure 1.**
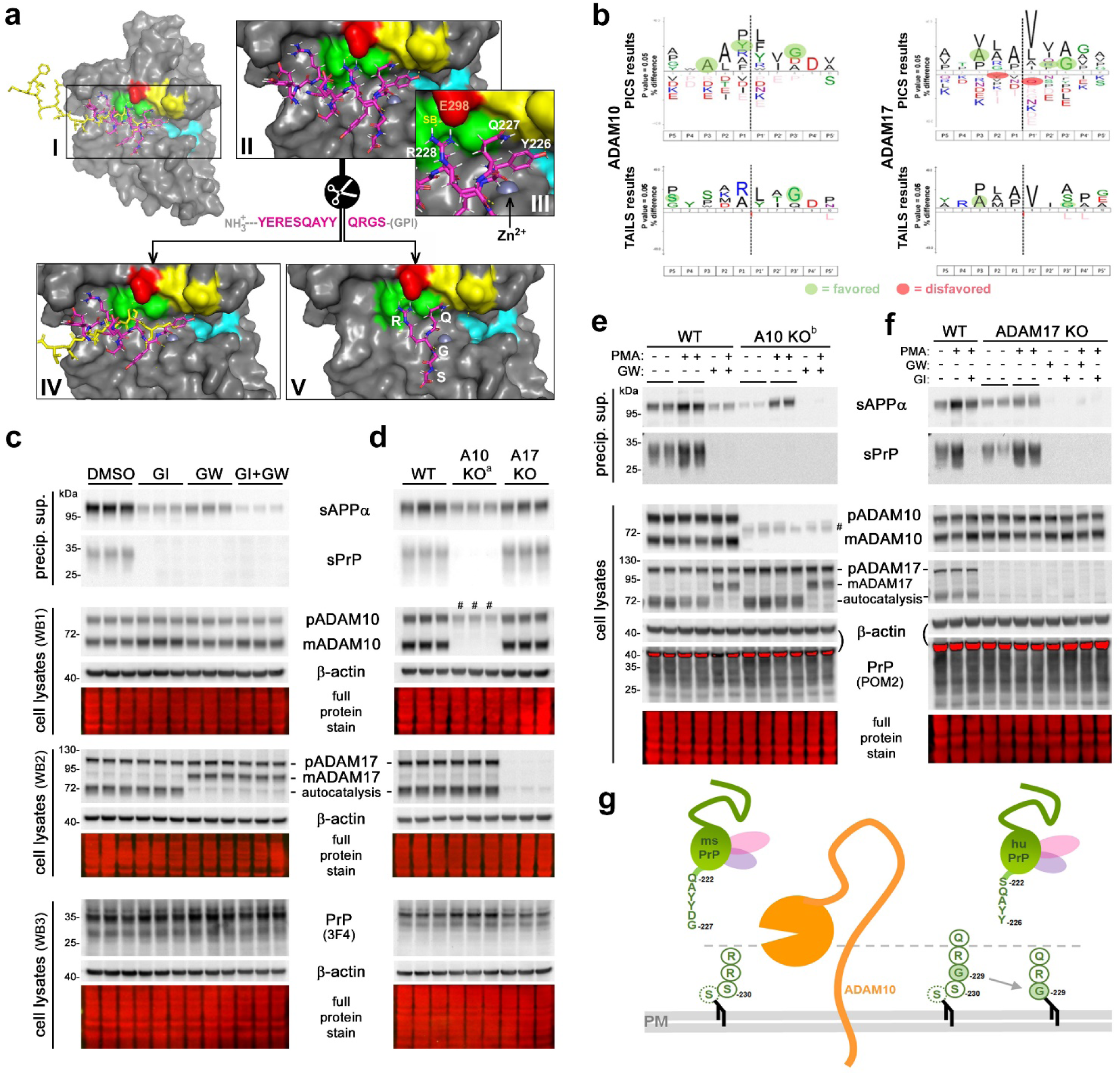
Cleavage site prediction based on structural models and pharmacological/genetic proof of PrP shedding in humans being ADAM10-dependent. (**a**) I: Overview of the proteolytic domain of human ADAM10 shown in surface representation (based on ^68^) with its divalent zinc ion (Zn^2+^; light grey ball) coordinated in the catalytic center. Key residues of substrate binding pockets are highlighted in yellow for S1 (V297, F323, D325, V327), cyan for S1’ (V376, I379, T380, I416, T422) and green for S3 (L301, L330, W332). Overlayed peptides of the extracellular C-terminal sequence ^642^FMRCRLVDADGPLG^655^ (yellow) of another ADAM10 molecule from the crystal structure (PDB: 6BE6) and the C-terminal end of human PrP ^217^YERESQAYYQRGS^230^ (purple) are shown as stick; the GPI anchor normally attached to the last serine is not depicted. Magnified view (II) and detailed selection (III) of the PrP C-terminal sequence positioned inside ADAM10’s catalytic domain indicating potential (transient) formation of a salt bridge (SB) between PrP R228 and ADAM10 E298 (red) and close proximity of the suspected cleavage site Y226↓Q227 and Zn^2+^ within the catalytic cave. IV: After cleavage, the N-terminal parts (C-termini of shed PrP in purple or soluble ADAM10 in yellow may be released. V: The remaining C-terminal residues of PrP may be freed from the catalytic domain (possibly regulated by the salt bridge) and stay attached to the membrane via their GPI anchor, from where they may be taken up for degradation. (**b**) IceLogos showing preferred and disfavored amino acids in different positions (P) to the potential cleavage site (P1↓P1’) based on various peptide and protein substrates of ADAM10 and ADAM17 using PICS (modified from ^71^) and TAILS (modified from ^72^). In case of putative PrP shedding, favored residues are highlighted by pale green, disfavored residues by pale red background. (**c**) Immunoblot analysis of sPrP and sAPPα (in TCA precipitated conditioned media) as well as PrP, ADAM10 and ADAM17 (in cell lysates) of A549 cells treated with metalloprotease inhibitors GI254023X (GI) or/and GW280264X (GW). β actin and total protein stain served as loading controls. (**d**) Assessment as before but in wild-type (WT), ADAM10 knockout (A10 KO) and ADAM17 knockout (A17 KO) A549 cells. (**e**) Biochemical analysis as before but in WT and A10 KO A549 cells treated or not with ADAM-stimulating phorbol ester PMA and/or inhibitor GW. Note that in **d** and **e** two different independently generated A10 KO A549 lines were used (A10 KO^a^ and A10 KO^b^) and both present with a different unspecific band/inactive mutant (marked in the blots with #). (**f**) Western blot analysis as before but in WT and A17 KO A549 cells treated or not with PMA and/or either inhibitor GW or GI. Red saturated bands in **e** and **f** result from residual β actin signal when reprobing for PrP as indicated. (**g**) Model of the membrane-proximate shedding of PrP. With the recent suggestion of glycine 229 (instead of previously assumed C-terminal serine) being the actual GPI anchor attachment site in human PrP ^85^, the same distance between cleavage site and membrane would be preserved between mice and humans. Correct size relations not depicted here to simplify matters.

Although neither us nor others ^22,23,67^ ever found indications of an involvement of the closely related ADAM17/TACE (with which ADAM10 shares several other substrates) in the C-terminal shedding of PrP, we also considered this metalloprotease and found some favored (A224 and G229 in PICS and TAILS, R228 in PICS) as well as disfavored PrP residues (Y225 and Q227 in PICS) (**Fig. 1b**).

Supported by these predictions and previous experience in raising cleavage site-specific antibodies for murine shed PrP ^24^, one of our groups generated antibodies against this putative shedding site, possibly enabling identification of extracellular PrP ending at Y226 as the physiologically relevant shed form (sPrP) in humans. Accordingly, rabbits were immunized with a respective peptide sequence and resulting polyclonal antibodies harvested and affinity-purified as described in the ‘materials & methods’ section.

### Confirming the ADAM10-dependency of PrP truncated at Y226 in human cell lines

Upon generation of polyclonal antibodies (termed sPrP^Y226^) directed against this assumed neo-C-terminus (as done before for the murine system ^24^), we first aimed at testing the banding pattern and ADAM10-dependency of immunoblot signals detected with this antibody. We also assessed any possible involvement of ADAM17/TACE in this supposed PrP shedding. We analyzed human lung carcinoma cells (A549) given their described decent expression levels of the relevant proteins ^76,77^. Molecular weight (MW) and glycoform pattern of bands detected with the sPrP^Y226^ antibody in conditioned media were in line with earlier findings on murine sPrP ^24^ (**Fig. 1c-f**). Treatment with two metalloprotease inhibitors, GI254023X (with its much higher potency towards ADAM10 than ADAM17) and GW280264X (basically inhibiting both proteases) ^78^, alone or in combination, resulted in a lack of sPrP signal in conditioned media (**Fig. 1c**). While PrP shedding was completely absent upon GI treatment, this inhibitor had no effect on ADAM17 activity as judged by a previously reported postlysis autocatalytic processing step (i.e., mature ADAM17 cleaving itself into a slightly shorter fragment upon cellular breakup ^79^) which was only inhibited in the presence of GW (**Fig. 1c**). In supposed contrast to PrP shedding, both proteases contribute to the non-amyloidogenic α-processing of APP as confirmed here by the differential influence of respective inhibitors on sAPPα levels in media supernatants ^80–83^. These results were also confirmed in a human brain-derived glioblastoma cell line (U373-MG; **Suppl. Fig. 1**). However, considering the known cross-inhibition and imperfect ‘specificity’ of the metalloprotease inhibitors for particular ADAM members (i.e., GW also inhibiting ADAM10 activity), we further investigated A549 cells depleted via CRISPR-Cas9 in ADAM10 (A10 KO) or ADAM17 (A17 KO). This analysis revealed that a knockout of ADAM17 had no effect on levels of sPrP, whereas shedding was completely abolished in the absence of catalytically active ADAM10 (**Fig. 1d**). To further exclude that ADAM17 may only participate in PrP shedding upon stimulation, we added the phorbol ester PMA, a widely used stimulus for ADAM17 activity, to WT and ADAM10 KO (**Fig. 1e**) or ADAM17 KO (**Fig. 1f**) A549 cells in the presence or absence of ADAM inhibitors. Although PMA treatment increased PrP shedding in WT cells, this effect was also observed in A17 KO cells, whereas no sPrP was detected in A10 KO cells. While this may support an influence of PMA on ADAM10 (e.g., via increased protein kinase C-mediated surface transport of ADAM10 ^84^), the stimulated PrP shedding is clearly independent from ADAM17 expression. In sum, this analysis confirmed the strict dependence of the immunoblot signal obtained with the new sPrP^Y226^ antibody on ADAM10, supporting that Y226↓Q227 might indeed be the relevant shedding site in humans. It further suggested sole involvement of ADAM10 in this shedding event, as described earlier for rodents ^22,23,67^.

Notably, membrane interaction of the catalytic domain of ADAM10 and distance of cleavage sites within ADAM10’s substrates to the plasma membrane are relevant aspects for shedding to occur ^74^. In this regard, it is intriguing that a recent study ^85^ suggested the GPI anchor in human PrP being attached to glycine 229 (instead of the subsequent serine residues as previously assumed (Source: www.uniprot.org; Major prion protein [Homo sapiens], ID: P04156)), which would preserve a similar distance between membrane and shedding site as in mice (**Fig. 1g**).

### Direct comparison of a poly- and a monoclonal antibody confirms PrP ending at Y226 as the product of ADAM10-mediated shedding in humans

Several years ago, one of our groups generated a set of mouse monoclonal antibodies against different C-terminally truncated forms of human PrP. Among those antibodies, one (termed V5B2) was described to specifically detect a shortened form of PrP ending at Y226 in the brains of a few patients suffering from prion disease ^61–64,86^. The fragment was then designated PrP226* ^87^ and appeared to accumulate in prion aggregates and to even correlate with the spatial distribution of PrP^Sc^ deposits ^65^. It was further characterized *in vitro* to predict structural and thermodynamic parameters affecting involvement in amyloid diseases ^88,89^. However, although the V5B2 antibody had been employed in several assays including ELISA, immunoblotting and immunohistochemistry (IHC), both the ‘mechanistic’ origin and physiological meaning of this fragment remained unclear until now, and there was no experimental support for it being a product of (physiological) ADAM10 proteolysis. We therefore directly compared the rabbit polyclonal sPrP^Y226^ antibody (introduced above) with the murine monoclonal V5B2 antibody. To this end, we first investigated the detection pattern of both antibodies in human neuroblastoma (SH-SY5Y) cells transiently transfected to overexpress human PrP (given the very low endogenous levels shown in the non-transfected control) (**Fig. 2a**). Two replica blots, both containing cell lysates and respective precipitated media supernatants, were first probed with either sPrP^Y226^ or V5B2 antibody. Both yielded very similar signals only in media samples of transfected cells yet not in respective cell lysates, and no signal was detected in media of cells treated with the ADAM10 inhibitor. When reprobed with the pan-PrP antibody POM2, strong overexpression of PrP in the lysates of transfected cells was confirmed. This overexpression may explain the lack of further elevated sPrP levels upon treatment with either Carbachol (a drug normally able to increase PrP shedding as shown in **Suppl. Fig. 2** and reported elsewhere ^25^) or PrP-directed IgGs known to stimulate shedding ^27^, as endogenous ADAM10 might simply be ‘saturated’ by the artificially high levels of PrP. In sum, both sPrP^Y226^ and V5B2 yield highly comparable WB signals, specifically detecting human ADAM10-cleaved shed PrP while being “blind” for its cell-associated full-length ‘precursor’.

**Figure 2.**
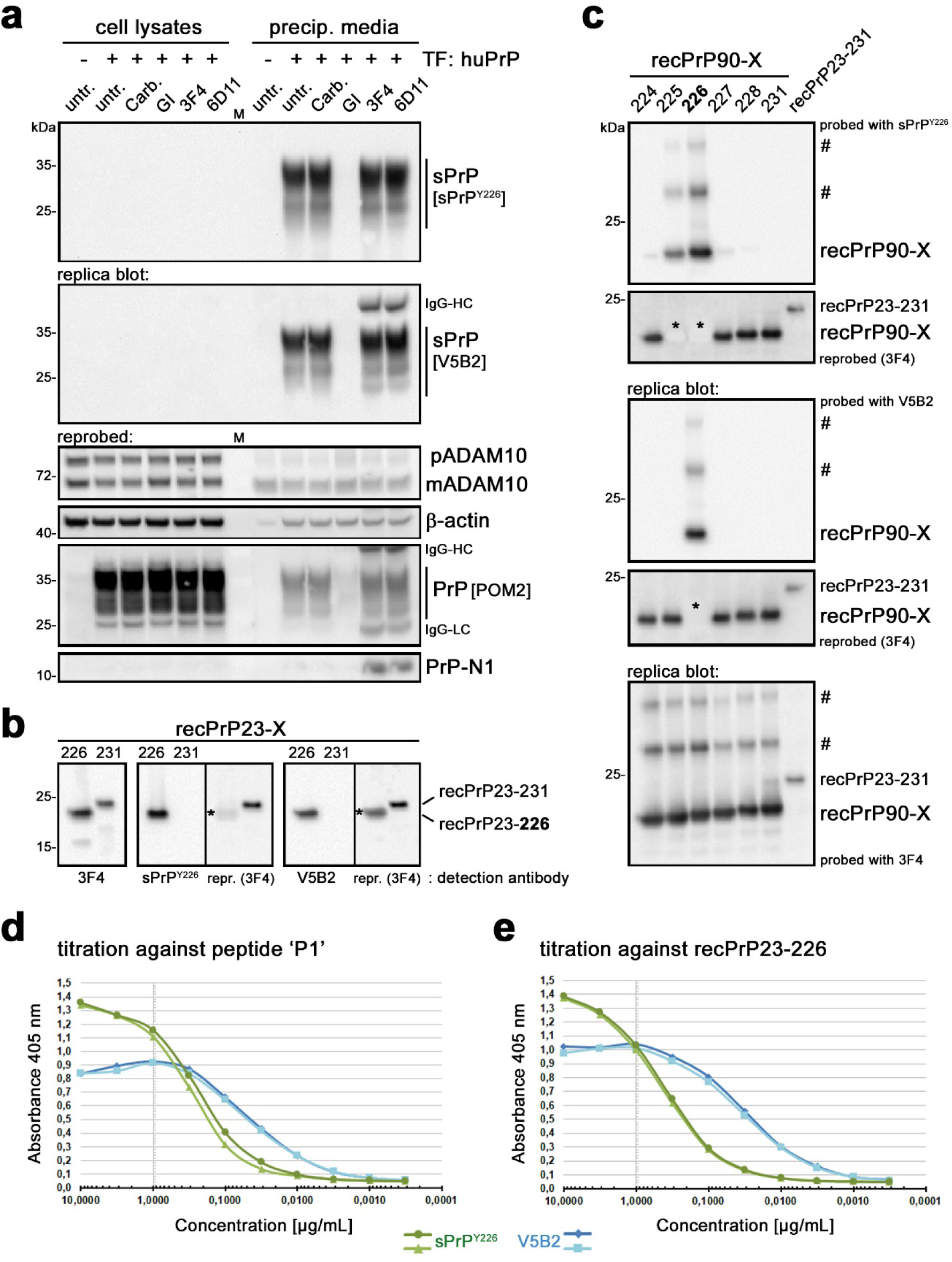
Direct comparison of polyclonal sPrP^Y226^ and monoclonal V5B2 antibodies. (**a**) Human neuroblastoma (SH-SY5Y) cells almost lacking endogenous PrP expression (see PrP signal in lysates of non-transfected (-) cells; left lane) were transfected (+) with a plasmid coding for human PrP and left untreated (untr.) or treated with carbachol (Carb.), ADAM10 inhibitor GI, or PrP-directed antibodies 3F4 or 6D11. Cell lysates (left half of blots) and precipitated media supernatants (right half) were loaded on two replica blots and initially detected with either sPrP^Y226^ or V5B2 yielding comparable signals (note that heavy chains (HC) of the murine treatment antibodies 3F4 and 6D11 are also detected with the anti-mouse secondary antibody used for detection of V5B2). Re-probing with the pan-PrP antibody POM2 confirmed strong overexpression of PrP in transfected cells (note that this cell-associated PrP was neither detected with sPrP^Y226^ nor with V5B2). Levels of premature (p) and mature/active (m) ADAM10, β-actin (loading control) and an N-terminal PrP fragment (N1) were also assessed. Actin signal in media likely indicates some degree of transfection-induced cell death (note the comparably weak signal in untransfected cells). Dominance of mADAM10 in media is likely associated with extracellular vesicles. Detectability of soluble PrP-N1 is rescued in the presence of 3F4 and 6D11 antibodies, which protect this instable fragment from C-terminal proteolytic trimming and degradation ^90^. ‘M’ indicates a protein marker lane. (**b** and **c**) Both antibodies, sPrP^Y226^ and V5B2, were also compared on replica immunoblots of recPrP23-226 (mimicking sPrP) versus recPrP23-231 (full length) (**b**) or of N-terminally (i.e., at aa 90) truncated recPrP with different C-termini (X, as indicated) as well as full-length recPrP (23-231) (**c**). Blots were re-probed and an additional replica blot directly detected with the pan-PrP antibody 3F4. Asterisks in re-probed blots in **b** and **c** mark “burned” signals resulting from overexposure during the previous detection shown above. ‘#’ indicate SDS-stable dimers or oligomers of the respective recPrP forms. (**d** and **e**) Comparison of sPrP^Y226^ (green curves) and V5B2 (blue curves) in ELISA against the peptide ‘P1’ initially used for immunization of mice to generate V5B2 (**d**) or against human recPrP23-226 (**e**)

Next, we analyzed the specificity of the antibodies against recombinant PrP variants ending at either position 231 or Y226. While a pan-PrP antibody detected both forms, polyclonal sPrPY226 and monoclonal V5B2 antibodies only detected the truncated recPrP ending at Y226 (**Fig. 2b**). We then wondered about the epitope tolerance of both antibodies and assessed their ability to detect different C-terminally truncated versions of human recPrP90-X by western blotting. For a cleavage site-specific monoclonal antibody, one would expect one exclusive signal for PrP90-226, whereas an analogue polyclonal antibody (due to its potential ‘repertoire’ of different IgGs) could conceivably also provide signals for fragments with neighboring C-terminal endings. Our analyses exactly confirmed this assumption as monoclonal V5B2 solely detected recPrP90-226, whereas polyclonal sPrP^Y226^ also detected few other fragments, albeit with much lower sensitivity (**Fig. 2c**). However, since such fragments likely do not exist in nature (despite the possibility of some rare stop mutations), the polyclonal sPrP^Y226^ antibody –like the monoclonal V5B2– can be regarded as a bona fide cleavage site-specific detection tool for human sPrP. Both antibodies also revealed bands at higher molecular weight likely representing SDS-stable oligomers of respective recombinant PrP variants.

To further examine sPrP binding propensity of V5B2 and sPrP^Y226^, we compared their relative binding affinities (RBA; i.e., concentration of the antibodies at half of the saturation, expressed in moles, M=mol/L) with ELISA. We performed titration experiments either against peptide ‘P1’ (CITQYERESQAYY, used for V5B2 generation ^61,62^) (**Fig. 2d**) or against recombinant human PrP ending at Y226 (recPrP23-226) (**Fig. 2e**). Despite slight differences in the curves, both antibodies showed a relatively high affinity with resulting RBAs against ‘P1’ of 2.0*10^-10^ (V5B2) and 1.0*10^-9^ (sPrP^Y226^) and against recPrP23-226 of 1.3*10^-10^ (V5B2) and 1.3*10^-9^ (sPrP^Y226^). This further supports their overall similar binding characteristics and usability in various methodological approaches. However, we also noted that the polyclonal antibody might be slightly better suited for detection of sPrP in denatured samples (as indicated by immunoblot comparison on serial dilutions of human brain; **Suppl. Fig. 3**), whereas its monoclonal pendant might be superior for native samples (as suggested by the ELISA results and a better performance in immunoprecipitating sPrP from conditioned cell culture media; **Suppl. Fig. 4**).

### Ligand-induced shedding of PrP in human cells

We have previously shown in the murine system that treatment of cells and brain slice cultures with certain PrP-directed antibodies stimulates the ADAM10-mediated shedding in a substrate-specific manner ^27^. Moreover, as shown above and earlier ^24^, shedding is completely abolished with an ADAM10 inhibitor. To investigate if these manipulations also work in the human neural system, and to further confirm that PrPY226 indeed corresponds to genuine, physiologically shed PrP, we employed these paradigms to three human brain-derived cancer cell lines which we had screened before for relevant endogenous expression of both ADAM10 and PrP. In our previous study using murine cells and tissues, 6D11 (an antibody binding to a central region in PrP) caused highest shedding among the PrP ligands tested, whereas the 3F4 antibody served as a negative control (as its epitope is absent in murine PrP yet present in human PrP). In the human cancer cell lines SHEP2 (neuroblastoma-derived; **Fig. 3a**), LN235 (astrocytoma-derived; **Fig. 3b**) and U373-MG (glioblastoma-derived; **Fig. 3c**), both antibodies –as expected– stimulated the shedding when compared to controls (albeit with only moderate effects of 6D11 in SHEP2 cells). Moreover, as shown before in mice, shedding of diglycosylated PrP seems to be preferred over the other glycoforms (as judged by comparison with the PrP glycopattern in respective lysates). In further agreement with murine samples and fitting to the lack of the GPI anchor and the very C-terminal amino acids, sPrP bands run at a slightly lower molecular weight than PrP in lysates. Besides a different ratio of premature and mature ADAM10 between the cancer cell lines (**Fig. 3a-c**), no obvious differences in PrP or ADAM10 levels were observed in cell lysates upon treatments.

**Figure 3.**
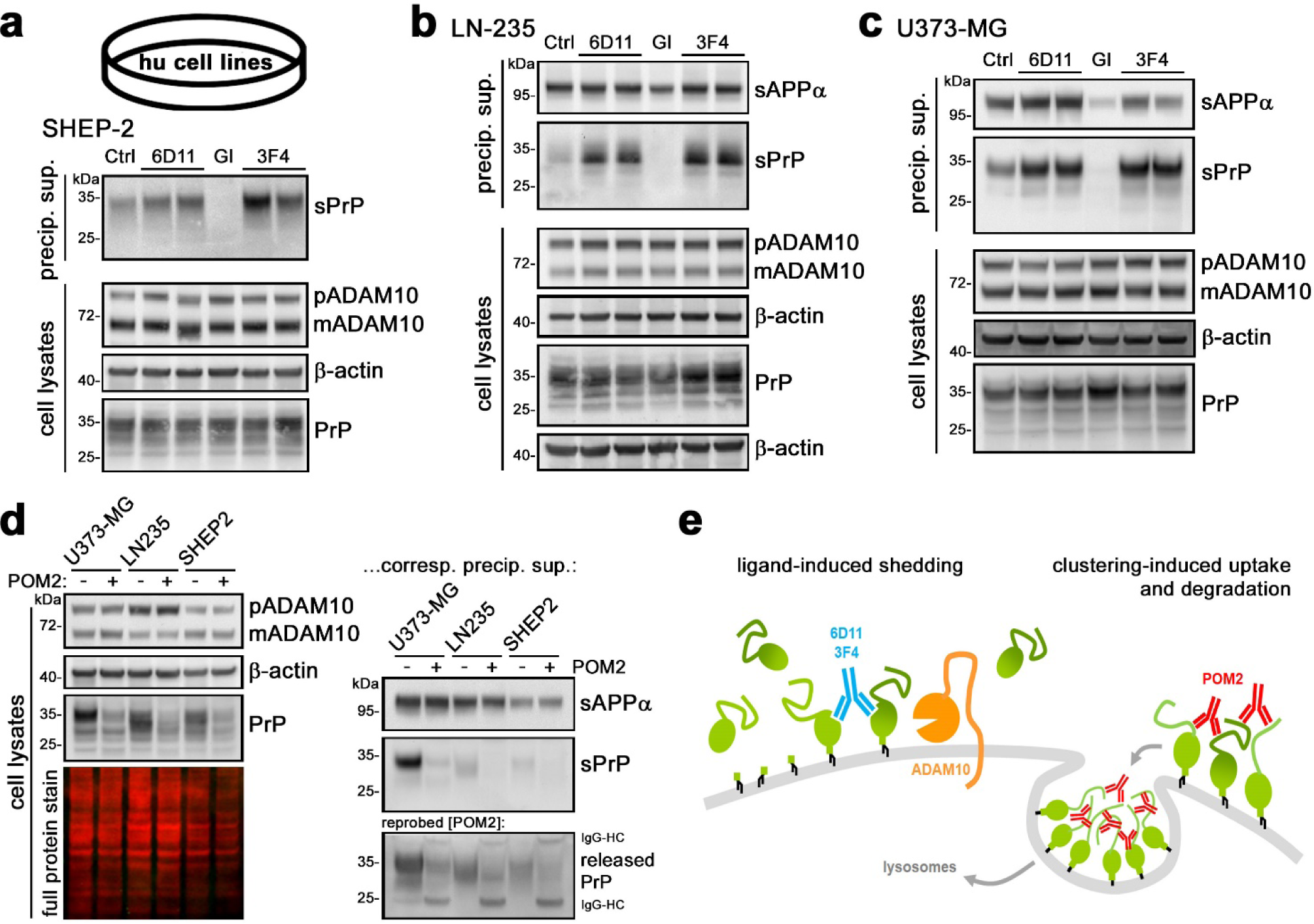
Proteolytic shedding of PrP, ADAM10 inhibition, and effects of PrP-directed antibodies in three human cancer cell lines of neural origin. Representative western blots showing basal levels of sPrP (Ctrl; left lane of each blot) detected with the sPrP^Y226^ antibody in precipitated overnight media supernatants of neuroblastoma cells (SHEP-2; in **a**), astrocytoma cells (LN-235; in **b**) and glioblastoma cells (U373-MG; in **c**). In all cell lines, shedding is increased upon treatment with PrP-directed antibodies 6D11 and 3F4 and abolished when treated with an ADAM10 inhibitor (GI). sAPPα was detected in media (in **b** and **c**) as another cleavage product generated by ADAM10. Corresponding cell lysates assessed for levels of PrP, premature (p) and mature/active (m) ADAM10, and β-actin (serving as loading control) are shown underneath (**a**-**c**). (**d**) Treatment with the antibody POM2 in all three cell lines results in the reduction of cell-associated PrP levels (left panel) as well as sPrP and released PrP in corresponding media samples (right panel). (**e**) Scheme showing the shedding-stimulating effect of PrP-directed antibodies and the exceptional reduction in total PrP levels caused by POM2 IgG (illustration modified from ^27^)

With regard to the shedding-stimulating effect of PrP-directed antibodies, our previous study on murine samples revealed one striking exception: POM2 IgG, an antibody directed against four repetitive epitopes along the flexible N-terminal tail of PrP, instead of increased shedding rather causes a general reduction in PrP levels in both cell lysates and corresponding media supernatants. This is due to clustering and multimerization at the cell surface triggering internalization and lysosomal degradation ^27^. Here, we also addressed this aspect and found the PrP-reducing effect of POM2 in the three human cell lines investigated (**Fig. 3d**).

Together with the complete inhibition of shedding with the ADAM10-specific inhibitor in all three cell lines of neural origin (**Fig. 3a-c**), these data strongly support both, (i) Y226 being the relevant cleavage site for ADAM10 in human PrP and (ii) our PrPY226-directed antibodies specifically detecting endogenously generated shed PrP. Moreover, the shedding-stimulating effect of PrP-directed antibodies as well as the downregulation of total PrP levels caused by POM2 IgG (both illustrated in **Fig. 3e**) are reported here for the first time in a human paradigm.

### Manipulation of PrP shedding in human neuronally differentiated embryonic stem cells and iPSC-derived brain organoids

Having confirmed in different human cell lines that PrP ending at Y226 is identical with physiological proteolytically shed PrP in humans, we directly assessed its presence and pattern in more complex cellular systems of human origin. Human neural stem cells (NSC) transduced to overexpress GFP-tagged PrP (due to low endogenous levels) were differentiated into a neuronal lineage (**Fig. 4a**). Cultures were then treated to manipulate the ADAM10-mediated shedding of PrP (as done before in cell lines). While treatment with the ADAM10 inhibitor impaired the shedding, PrP-directed antibodies 3F4 and 6D11 caused increased levels of sPrP in conditioned media (**Fig. 4b**). We next investigated this in human iPSC-derived cerebral organoids (**Fig. 4c**). After 5 months of culture, expression levels of ADAM10 and PrP were good, sPrP was detectable in conditioned media (**Fig. 4d**), and diverse brain cell types (except for microglia) constituted the organoid as confirmed by expression of typical markers (**Fig. 4e**). Again, shedding of PrP was abolished upon GI-treatment, whereas it was increased upon incubation of cells with the 3F4 antibody binding to human PrP (**Fig. 4f**).

**Figure 4.**
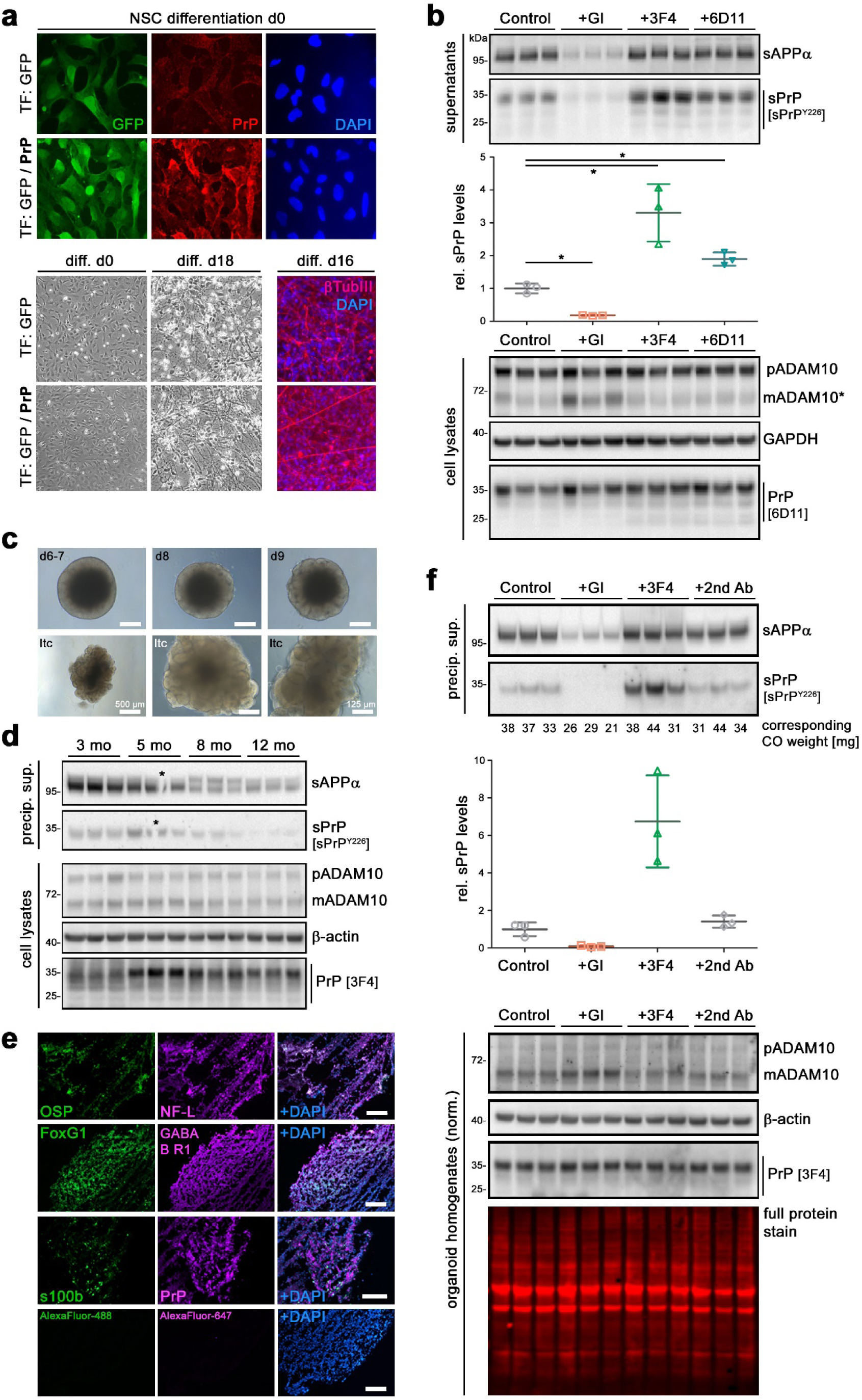
PrP shedding in human neuronally differentiated stem cells and iPSC-derived cerebral organoids. (**a**) Immunofluorescence (IF) analysis of human embryonic stem cell-derived NSC (upon lentiviral transfection to express either GFP (green) alone or GFP and exogenous PrP (red)) at day 0 of neuronal differentiation (upper panel). Bright field microscopy (lower left panel) showing morphological differences between day 0 and 18 of neuronal differentiation. IF analysis at day 16 (lower right panel) reveals expression of neuronal marker β-tubulin III. (**b**) Immunoblot assessment of sPrP and sAPPα in conditioned media (supernatants; upper panel), densitometric quantification of relative sPrP levels (diagram in middle panel), and cellular levels of ADAM10, GAPDH (as loading control) and PrP (lysates; lower panel) in NSC following 30 days of neuronal differentiation and 18 h of treatment with ADAM10 inhibitor GI (note that a lower concentration [6µM] was used here, hence the residual signal for sPrP) or PrP-directed IgGs 3F4 or 6D11. Controls treated with DMSO (as in GI-treated samples) served as reference (set to 1 in quantification). Quantification: n=3 wells per condition; mean ± SE; Student’s *t*-test with \**p* < 0.05. (**c**) Human iPSC-derived cerebral organoids (CO) at different days (d) of differentiation and after neuroepithelial bud expansion ready for long-term culture (ltc). Scale bar = 250 µm unless otherwise indicated. (**d**) Levels of sPrP and sAPP (in conditioned media) and PrP, ADAM10 and β-actin (loading control) in homogenates of organoids after different culture durations (3-12 months). (**e**) Presence of different cell types within differentiated organoids confirmed by IF analysis of typical markers (OSP = oligodendrocyte-specific protein; NF-L = neurofilament light chain (for neurons); GABA B R1 = gamma-aminobutyric acid type B receptor subunit 1 (for inhibitory neurons); s100b = S100 calcium-binding protein B (for astrocytes)). PrP expression was also detected. DAPI was used to stain nuclei. Staining controls with only fluorescently labeled 2nd antibodies (AlexaFluor-488 or −647) revealed no signals. Scale bars = 200 µm. (**f**) Treatment of CO with GI (shedding inhibition), 3F4 antibody (shedding stimulation) and a non-specific secondary antibody (as negative control). Amounts of sPrP and sAPP in precipitated conditioned media (densitometric quantification of sPrP shown below) and ADAM10 and PrP in homogenates of respective CO (COs vary in size, which may influence the amount of proteins shed into the media, therefore the weights of individual COs are shown below the respective lane) assessed by immunoblotting. Full protein stain and β-actin shown as loading controls. Note that we refrained from statistical analysis given the variation in CO weights

### Heterologous cleavage occurs, and the new cleavage site-specific antibodies also detect shed PrP of animal species susceptible to naturally occurring prion diseases

Given the difference in C-terminal sequence and shedding sites between human and rodent PrP, we wondered whether heterologous cleavage (i.e., human ADAM10 shedding mouse PrP and vice versa) is possible. To this end, we first expressed human PrP in murine PrP-depleted N2a cells ^90^. Replica blots of the same conditioned media were probed either with our polyclonal antibody for rodent sPrP (sPrP^G227^) or with the respective counterpart for human sPrP (sPrP^Y226^) (**Fig. 5a**). A similar experiment was done including GFP-tagged versions of both species’ PrPs (**Fig. 5b**). Shedding occurred in all instances, indicative of ADAM10 being tolerant to other species’ sequences and preserving those PrPs’ cleavage sites. Similar results were obtained when expressing murine PrP in human SH-SY5Y cells (**Fig. 5c**). These experiments also further support the specificity of the different sPrP-directed antibodies. Moreover, detection of N1 (and N3) (in **Fig. 5a** and **5c**) and C1 fragments (in **Fig. 5b**) indicates that α-cleavage (and γ-cleavage), for which responsible proteases are not yet identified without doubt, also occur in a heterologous setting. Next, we assessed whether antibody sPrP^Y226^ raised against human sPrP would also detect sPrP in other species. Upon initial analyses of CNS samples from a broad range of domestic and zoo animals, we got a glimpse of some promising fragments (fitting to either sPrP or the shed N-terminally truncated C1 fragment) not only in human and macaque, but also in goat, sheep, cattle and two deer species (**Suppl. Fig. 5**). Fittingly, the latter species, in contrast to mice and rats, largely share the sequence around the cleavage site in human PrP (**Fig. 5d**). This prompted us to perform further analysis in transgenic mice (depleted for endogenous PrP) expressing PrP from either sheep, goat, cattle or human PrP (the latter with MM or VV status at polymorphic position 129). Using the sPrP^Y226^ antibody, sPrP forms were detected in the brains of all transgenic mice (**Fig. 5e**). This not only confirms the heterologous cleavage discussed above, but also reveals that PrP shedding in those major species prone to naturally occurring prion diseases (i.e., scrapie in sheep and goats; BSE in cattle, and CWD in deer) occurs after the respective tyrosin corresponding to Y226 in the human sequence. Here again, we directly compared the performance of both sPrP antibodies in some of these transgenic mice (versus WT and PrP-KO mice) by immunoblotting (**Suppl. Fig. 6**). This analysis supported similar overall detection profiles, yet also confirmed that the polyclonal version reveals stronger signals when detecting denatured samples.

**Figure 5.**
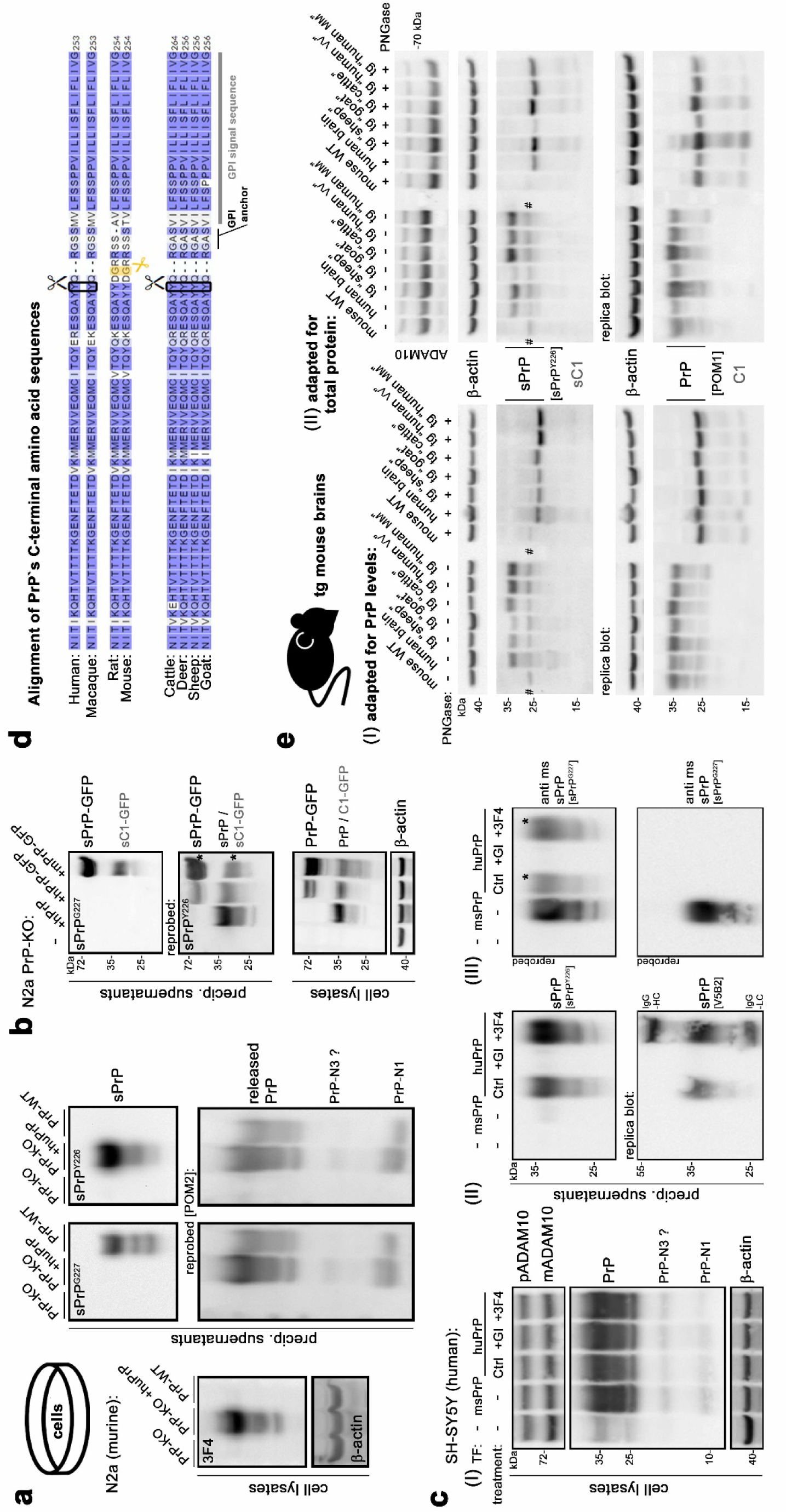
Heterologous cleavage and species-specificity of sPrP-directed antibodies. (**a**) Assessment of PrP shedding in murine PrP-KO N2a cells overexpressing human PrP (PrP-KO and WT N2a cells served as controls for no and endogenously expressed murine PrP, respectively). Overexpression of human PrP confirmed by a 3F4-positive signal in western blot of lysates. Replica blots of precipitated conditioned media detected with either sPrP^G227^ (for murine sPrP) or sPrP^Y226^ (for human sPrP). Presence of all released PrP fragments was confirmed by re-probing with POM2 antibody. (**b**) Western blot analyses of PrP-KO N2a cells transfected with human PrP or GFP-tagged versions of human or mouse PrP. Detection with sPrP^G227^ exclusively detects murine sPrP-GFP and shed C1-GFP, whereas re-probing with sPrPY226 reveals human sPrP, sPrP-GFP and shed C1-GFP in respective media samples. Expression of respective cell-associated PrP forms in lysates (using pan-PrP antibody POM1) is shown below. (**c**) Western blot analyses of human SH-SY5Y cells transfected (TF) with murine or human PrP (the latter treated or not with GI or 3F4 IgG). (I) Cell lysates; (II) replica blots of precipitated media detected with either polyclonal sPrP^Y226^ (top) or monoclonal V5B2 (bottom) detecting human sPrP (basal (Ctrl), inhibited (GI) or increased (3F4)); (III) re-probing with sPrP^G227^ reveals murine sPrP (* indicates previous signals resulting from the initial detection (II) due to primary/secondary antibody species combination). (**d**) C-terminal amino acid sequences of PrP in different species including GPI-anchor signal sequence and attachment site. The ADAM10 cleavage site is marked in yellow for rats and mice, in black for human and monkey PrP. Note the sequence similarity of the latter with cattle, deer, sheep and goat. (**e**) Assessment of sPrP and PrP in brain homogenates of transgenic (tg) mice expressing PrP of different species as indicated. A wild-type mouse brain and a human brain homogenate were included as controls. PNGase F digestion was performed for deglycosylation (shown on the right side of each blot). Protein amounts were either roughly adapted to PrP expression (I) or normalized for total protein (II). Actin served as loading control, and ADAM10 levels are also shown in II. # indicates presence of an unspecific band detected with sPrP^Y226^

### Altered distribution of sPrP with accumulation in extracellular deposits uncovers presence of PrP^Sc^ aggregates in different human and animal prion diseases

Some cases of human prion diseases are linked to *PRNP* stop mutations causing expression of pathogenic C-terminally truncated (and hence anchorless) PrP versions, which form large extracellular and often vessel-associated deposits. Notably, normal (i.e., non-genetically truncated) PrP expressed by the unaffected allele associates with these aggregates ^91–94^. But why and how should regular membrane-anchored PrP actually end up in plaques to a relevant extent? Since we recently showed in prion-infected mice that sPrP co-localizes with bona fide PrP^Sc^ deposits ^27^, we considered the aforementioned findings in patients may either reflect passive recruitment of sPrP to PrP^Sc^ deposits or even an active involvement of the shed form in extracellular sequestration of harmful oligomeric PrP^Sc^ assemblies. To address whether this interaction might be a more widespread phenomenon, we performed morphological assessment to directly assess tissue distribution of sPrP and its potential association with prion deposits in brain samples of human and –given the detection characteristics of our antibodies– animal prion diseases. As shown in **Fig. 6a**, in a control brain not diagnosed with neurodegeneration, sPrP appears uniformly like a background staining due to its diffuse and even distribution. In prion diseases, however, sPrP (using polyclonal sPrP^Y226^) becomes visible and, even without a harsh pre-treatment required for detection of bona fide resistant PrP^Sc^, indicates presence of extracellular prion aggregates both in sporadic CJD (sCJD) cases of the MM2C type (coarse-grained and perivacuolar deposits in the cortex) and the MV2K type (Kuru-like plaques in cerebellum), reminiscent of our earlier findings in mice ^27^. In cerebellum of the MV2K case, sPrP clusters were even more pronounced than actual PrP^Sc^ plaques, which may suggest a role as an aggregation hub for oligomeric misfolded proteins. Re-localization and aggregate association of sPrP was also observed when monoclonal V5B2 was used to stain brains affected by sCJD or variant CJD (**Fig. 6b**). A direct comparison of both antibodies (**Suppl. Fig. 7**) confirms a comparable detection pattern of sPrP^Y226^ and V5B2 in different sCJD subtypes and brain regions. Since cattle and sheep share the cleavage site (**Fig. 5**), we stained brain samples of these species affected or not by classical BSE and classical scrapie, respectively, and again found the sPrP-characteristic re-distribution in the presence of prion deposits (**Fig. 6c**). Immunofluorescence analysis of different human prion diseases (vCJD and Gerstmann-Sträussler-Scheinker (GSS) syndrome in **Fig. 6d** and sCJD in **Fig. 6e**) further revealed the intimate association of sPrP with prion plaques. Lastly, since heterologous shedding occurs (**Fig. 5**), we assessed transgenic mice expressing either ovine PrP (tg338, infected or not with NPU1 prions; **Fig. 6f** and **Suppl. Fig. 8a,b**) or bovine PrP (with or without vCJD infection; **Fig. 6f** and **Suppl. Fig. 8c**). Besides confirming the aforementioned partial colocalization of sPrP with the respective strain-characteristic extracellular prion deposits in different brain regions, we also found a pronounced vessel-associated pattern.

**Figure 6.**
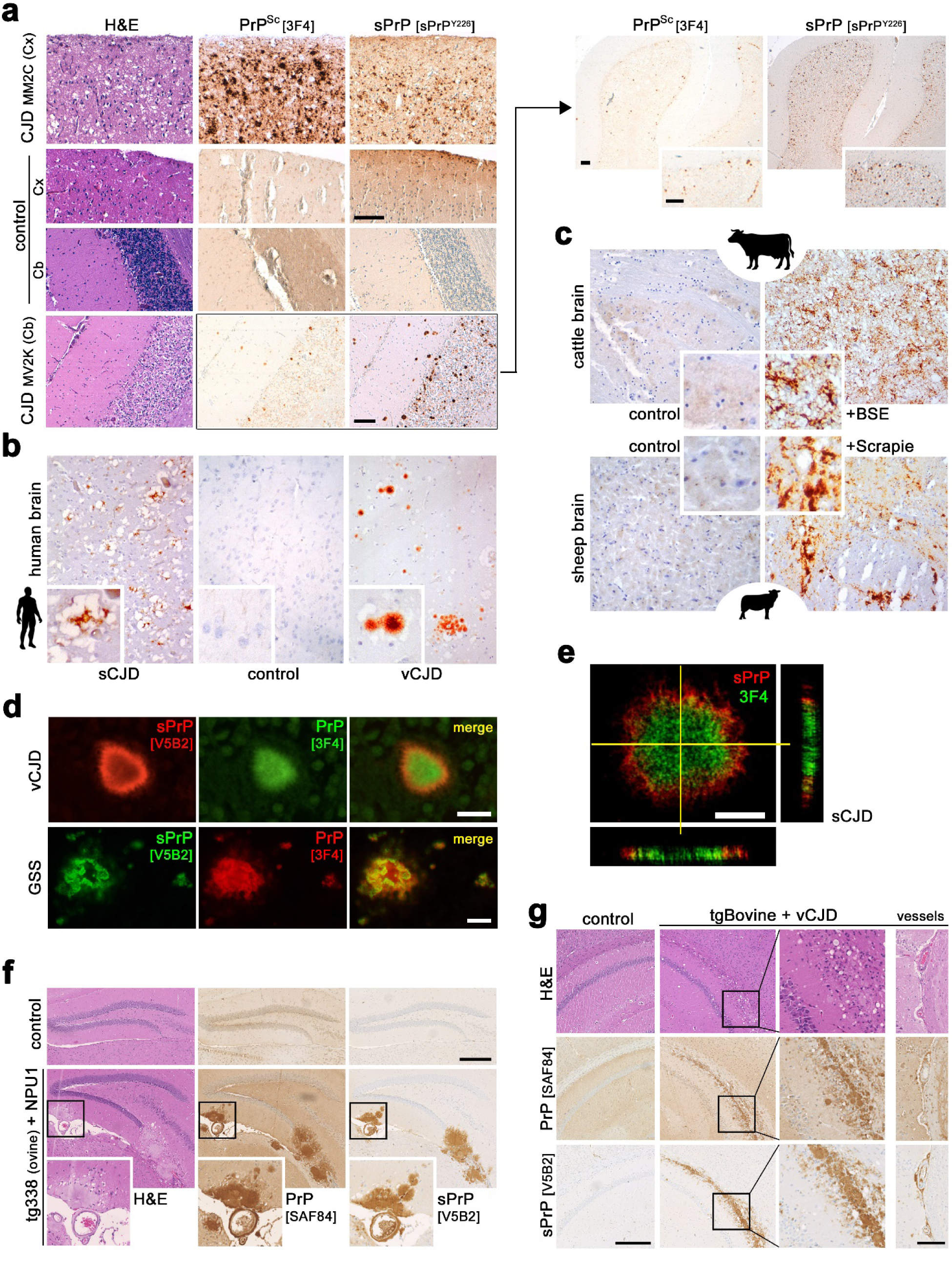
Redistribution of sPrP and association with prion deposits in prion diseases of humans and animals. (**a**) (Immuno)histochemical (IHC) assessment of PrP^Sc^ (3F4 antibody upon harsh tissue pre-treatment) and sPrP (polycl. antibody sPrP^Y226^) in two CJD cases compared to a control without diagnosed neurodegeneration. Coarse-grained and perivacuolar PrP^Sc^ deposits are seen in frontal cortex [Cx] of a MM2C case, while the cerebellum [Cb] of a MV2K case presents with typical Kuru-like plaques (note that tissue disruption in the control is due to the pre-treatment prior to PrP^Sc^ detection). Shed PrP shows a diffuse distribution in the control and re-distributes into an aggregated appearance in brains affected by CJD (scale bars: 100 µm). In the MV2K, more sPrP clusters appear than actual PrP^Sc^ plaques, which is further supported by an overview comparison (upper right panel). (**b**) Monocl. antibody V5B2 used in IHC to detect sPrP in brain sections of a sCJD and a vCJD patient (compared to a control). (**c**) Detection of sPrP (V5B2) in the brains of cattle affected or not with BSE (upper panel) and sheep with or without Scrapie (lower panel). (**d**,**e**) Immunofluorescence analyses showing association of sPrP (V5B2) with extracellular PrP aggregates (3F4) in acquired (vCJD) and genetic prion diseases (GSS) (**d**; standard fluorescence microscopy; scale bars: 20 µm) and sCJD (**e**; z-stacks with side projections; scale bar: 5 µm). (**f**,**g**) Histological analyses of large PrP^Sc^ deposits (here: SAF84 antibody without harsh pre-treatment) and sPrP (V5B2) in hippocampal areas of prion-infected transgenic mice expressing ovine (tg338; **f**) or bovine PrP (**g**). Tg338 mice infected with NPU1 prions present with large and dense amyloid-like plaques (**f**). TgBov mice infected with vCJD show extended prion deposition along the corpus callosum. Boxes indicate position of magnified areas (**g**). In both models, association of aggregates with brain vessels is observed. Non-infected mice of the respective genotype served as controls. Scale bars: 250 µm (and 100 µm for the ‘vessels’ panel in **g**)

### Shed PrP also closely associates with extracellular amyloid deposits in AD and is readily detectable in human CSF

In AD and other neurodegenerative diseases, large deposits of disease-associated misfolded proteins may be less harmful than their diffusible synapto- and neurotoxic oligomeric states ^95–99^. Earlier studies reported presence of PrP within Aβ deposits in AD brain ^15,100–102^, yet the mechanistic origin of this plaque-associated PrP remained obscure until our recent demonstration in mouse models that this particularly identifies as sPrP ^26,27^. This finding, together with the capacity of soluble PrP fragments to bind and detoxify Aβ ^30–33^ and the known ability of PrP to foster Aβ aggregation ^103,104^, suggest a protective sequestration of Aβ (and possibly other harmful PrP-binding extracellular oligomers alike) driven by sPrP. When analyzing sPrP levels in AD brain at different disease stages by western blot, we found interindividual differences yet no significant alterations between groups (**Fig. 7a**), fitting to similar total amounts of sPrP detected earlier in brains of 5xFAD mice and controls ^27^. Upon immunohistochemical assessment, however, we observed a marked redistribution of sPrP into structures reminiscent of larger diffuse deposits or smaller dense plaques of Aβ, which was absent in non-AD controls (**Fig. 7b**). Occasionally, we also found dense deposits associated with vessels in brains of patients with AD and those without diagnosed neurodegenerative disease. Immunofluorescence co-stainings in AD samples then revealed enrichment of sPrP in amyloid plaques, as seen before in mouse models for Aβ pathology (**Fig. 7c**) ^26,27^. When isolating microvessels from AD brain, in some instances extracellular plaques were co-purified and showed an intimate association between Aβ and sPrP (**Fig. 7d**). Moreover, this analysis also confirmed presence of sPrP in vessel-associated amyloid deposits (**Fig. 7d** and **7e**; orthogonal views shown in **Suppl. Fig. 9**). Lastly, we addressed detectability of sPrP in human CSF. When adjusted to balance total protein content, both sPrP and shed C1 (sC1; resulting from PrP α-cleavage followed by ADAM10-mediated shedding) were much more abundant in CSF than in brain homogenates (**Fig. 7f**), fitting to a soluble factor being drained into body fluids ^53^.

**Figure 7.**
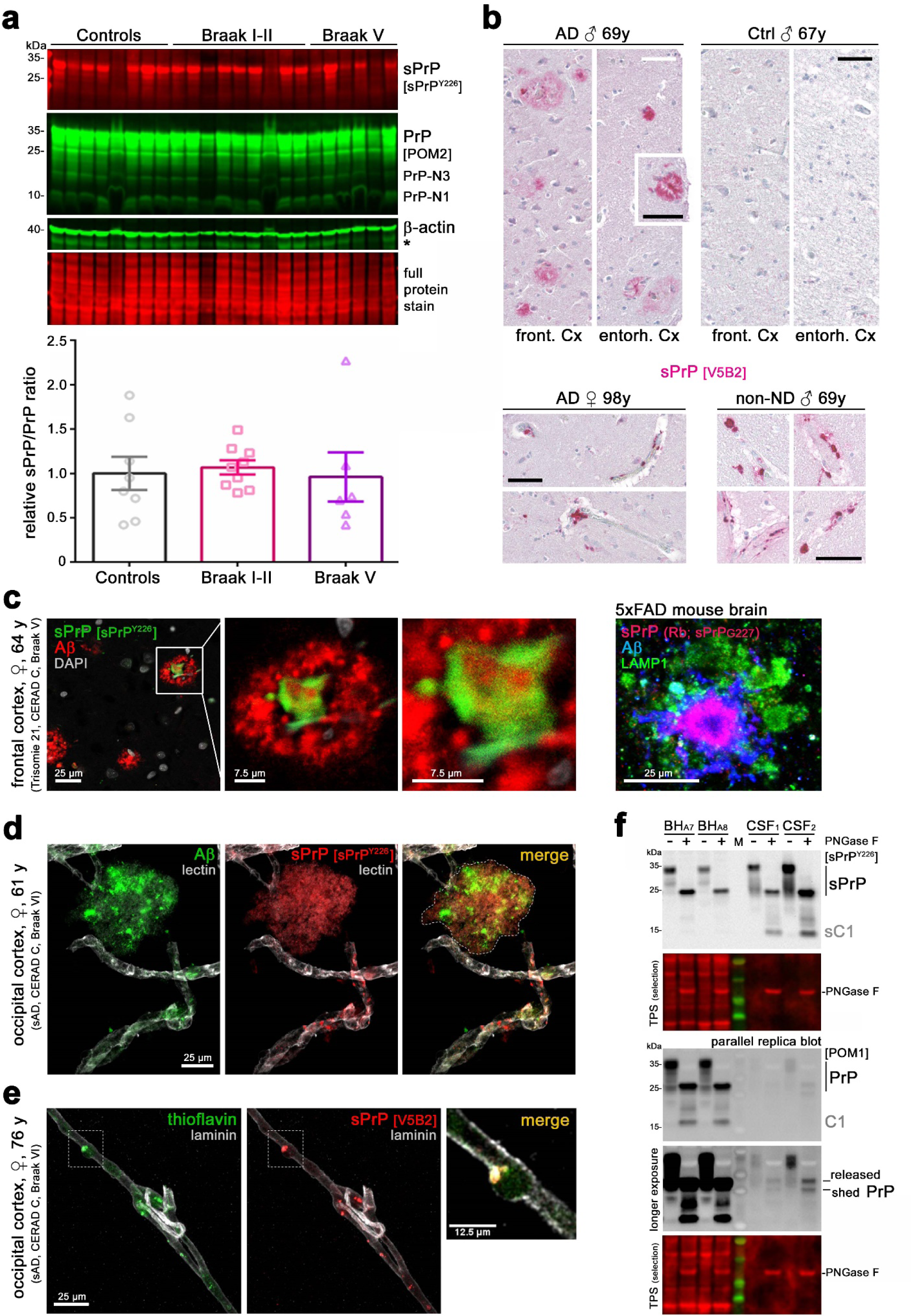
Analysis of shed PrP in AD and CAA and sensitive detectability of sPrP in human CSF. (**a**) Western blot analysis of sPrP and total PrP in frontal cortex samples of AD patients at Braak stage I-II (n=8) and Braak stage V (n=9) compared to non-neurodegeneration controls (n=6). Actin and total protein stain served as loading control. Asterisk indicates residual signal from previous PrP detection. Besides apparent inter-individual differences in sPrP levels, quantification (below) of the sPrP/PrP ratio did not reveal major (significant) differences between groups. (**b**) IHC assessment of sPrP in AD and control brains. In AD brain, sPrP shows both diffuse and dense (birefringent) plaque-like pattern reminiscent of bona fide amyloid-β (Aβ) deposits (upper panel). Dense vessel-associated sPrP signal can be found in some AD cases and controls (lower panel). Scale bars: 50 µm. (**c**) Closer inspection by immunofluorescence microscopy in brain sections of a patient with Trisomie 21 and AD reveals presence of sPrP in the center of some (yet not all) Aβ plaques, as seen earlier in respective mouse models ^27^ (on the right: staining for sPrP in this 5xFAD mouse brain was with sPrP^G227^ antibody; LAMP1 indicates dystrophic neurites or microglial lysosomes). (**d**) Large plaque-like clusters (highlighted by dotted line in the merge picture) of Aβ and sPrP were co-purified during isolation of microvessels from AD brain. Co-localization of both molecules was also found at/in vessels. Lectin was stained as endothelial marker. An orthogonal projection of z-stacks of this picture is presented in Suppl. Fig. 9a). (**e**) Association of amyloid and sPrP in/at brain vessels was verified in another AD case using another set of antibodies/stain (monocl. V5B2 for sPrP, thioflavin for (Aβ) aggregates, anti-laminin as endothelial/vessel marker). Another vessel of this sample is shown in orthogonal view in Suppl. Fig. 9b). Scale bars in c, d, e as indicated. (**f**) Immunoblot analysis of sPrP and total PrP in two human brain homogenates (BH) and two CSF samples (patients not diagnosed with neurodegeneration). Deglycosylation (+ PNGase F) was performed for better detection of the (shed) C1 fragment (resulting from ADAM10-mediated shedding after PrP α-cleavage). Note that 20 µg of total protein were loaded for BH, whereas CSF samples had only 1 or 3 µg of total protein. TPS=total protein stain

Thus, sPrP closely interacts with aggregating proteins associated with human neurodegenerative diseases. Since sPrP seems to be immobilized inside deposits of misfolded proteins and may hence be kept inside the brain in respective pathologies rather than being physiologically drained into the CSF, further studies addressing conceivable disease-related alterations in sPrP levels in body fluids are warranted regarding a diagnostic potential.

## Discussion

New pathomechanistic insight and potential therapeutic targets together with earlier diagnosis are urgently required in the field of currently incurable neurodegenerative diseases, ranging from rather rare transmissible prion diseases to Alzheimer’s disease, the most frequent cause of dementia. Focusing on the proteolytic processing of PrP, a common denominator in these conditions of the brain ^6–16^ and potentially relevant player in other pathophysiological processes throughout the body ^26,45–48,50–52^, we here formally demonstrate that ADAM10 is the physiological sheddase of PrP in humans, mediating the release of nearly full-length PrP from the plasma membrane. We identified the cleavage site in humans (and some mammalian species most relevant for natural animal prion diseases) and present cleavage site-specific antibodies allowing to detect sPrP with a variety of techniques and in different biological samples. We also provide the first demonstration that human brain sPrP, usually diffusely distributed in the extracellular space, is relocalized in the presence of extracellular deposits of misfolded proteins, closely associating with the latter. While this may support a protective sequestrating activity of sPrP towards toxic protein assemblies, it confirms earlier findings in mouse models ^26,27^ and provides a mechanistic rationale for the widespread earlier observation of “normal PrP” being enriched in extracellular protein deposits in diverse human proteinopathies ^15,91–94,100–102,105^. A scheme summarizing the key findings is provided in **Fig. 8**.

**Figure 8.**
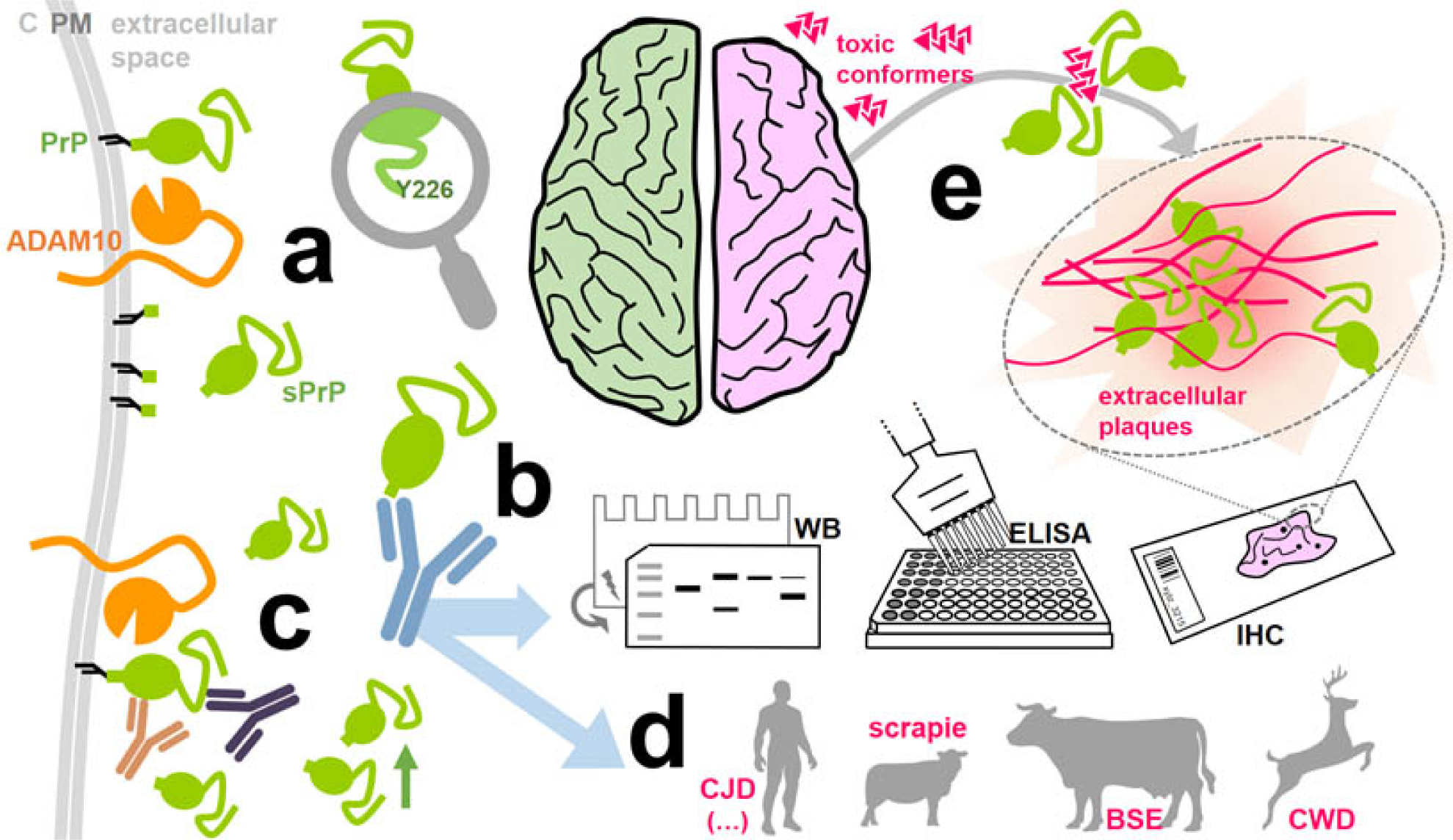
Graphical summary. (**a**) The widely expressed metalloprotease ADAM10 (orange) is the functionally relevant sheddase of PrP (green) in the human body and constitutively releases shed PrP (sPrP) into the extracellular space, from where it is also drained into body fluids such as CSF (not depicted to simplify matters). C = cytoplasm; PM = plasma membrane. We here identified the cleavage site between PrP’s tyrosine 226 and glutamine 227. (**b**) We generated cleavage site-specific antibodies against this neo-C-terminus (Y226). The sPrP-specific poly- and monoclonal antibodies do not detect full-length membrane-bound forms of PrP and can now be used in several routine methods, such as western blotting (WB), ELISA, and immunohistochemistry (IHC), to analyse a wide range of biological samples in basic science and diagnostics. (**c**) As shown before in mice, we demonstrate that PrP shedding can also be stimulated in the human system by PrP-directed ligands (e.g., antibodies), a mechanism of potential therapeutic value. (**d**) We also found that the cleavage site in human PrP is shared by other mammals including sheep/goats, cattle and deer. Hence, the sPrP-specific antibodies presented here will also foster analyses in the most relevant species (naturally) affected by prion diseases. (**e**) Among other findings, we show that sPrP redistributes from a diffuse pattern (in healthy brain) to markedly cluster with extracellular deposits of misfolded proteins in neurodegenerative diseases of humans and animals, possibly pointing towards a protective sequestrating activity of sPrP (containing all relevant binding sites) against toxic diffusible conformers in the extracellular space

The interest in endogenous proteolytically generated PrP fragments is steadily increasing ^19,20,106–109^ with more and more functions and pathological implications being suggested, particularly for ‘sPrP’ ^45–51^. Yet due to the lack of appropriate tools, most studies either did not sufficiently discriminate between sPrP and other released PrP forms (e.g., on EVs) or drew their conclusions from experiments using synthetic PrP, considering the latter to be a suitable analogue for physiological sPrP. This may or may not be the case (given the potentially relevant differences in glycosylation state and C-terminal ending ^26^). Cleavage site-specific antibodies, now available for rodents ^24^ and the human system (as presented here), will certainly be valuable in clarifying these and future questions.

With regard to neurodegenerative diseases, promising PrP-related therapeutic strategies include reduction of total or cell surface PrP levels (e.g., via antisense oligonucleotides (ASOs) ^110–112^ or other compounds ^113^) and treatment with PrP-directed antibodies or other ligands (aiming to block membrane-bound PrP’s interaction with toxic conformers and/or to stabilize its native fold ^114^ (reviewed in ^27^)). Our previous identification in murine samples of a ligand-stimulated shedding of PrP ^27^ adds another mode of action, linking both concepts and likely contributing to protective effects ascribed to PrP-binding antibodies. Enabled by our sPrP-specific antibodies used for detection, we here show that this mechanism also applies to the human context. Though speculative at the moment, combination therapies are conceivable. A PrP expression-lowering approach, for instance, could be combined with stimulated shedding to “transform” the remaining (harmful) membrane-bound PrP into a released (protective) anchorless factor (while possibly even preserving physiological ligand functions of sPrP).

PrP levels in body fluids not only serve as disease biomarker ^115,116^, but also as a surrogate marker for treatment efficacy, e.g. in ASO-based PrP-lowering strategies ^117,118^. However, “PrP” in this context rather represents a pool of different iso-and proteoforms ^5,19^ and, in addition, is enriched on certain EV subtypes ^24,119^. A compensatory network connects mechanisms of cellular PrP processing and release ^24^, yet how production of the different PrP forms is regulated and how it would react to manipulation of PrP expression is unknown. Available pan-PrP antibodies, depending on their epitopes, would either not discriminate between diverse differentially regulated and affected PrP subforms or could be unresponsive for some of the latter. Reliable detection of a well-defined fragment, such as sPrP, and treatment-associated alterations therein could therefore be superior, highlighting the likely diagnostic potential of the cleavage site-specific antibodies presented here ^117,118,120^.

Reliable surrogate markers are critical when it comes to pharmacological targeting of highly disease-relevant enzymes such as secretases ^121–123^. The rather ubiquitous expression of both ADAM10 and PrP in different organs, cell types and experimental models, and the fact that no other proteases (such as ADAM17) seem to be involved in PrP shedding, highlight the potential of measuring sPrP as a surrogate marker for efficacy read-out in any experimental or therapeutic strategies targeting ADAM10, be it stimulation of its protective APP α-secretase activity in the context of AD or inhibition of its rather detrimental effects, e.g. in cancer and inflammatory diseases ^124^. Hence, future studies aiming to manipulate ADAM10 may take advantage of sPrP-specific antibodies in basic research and clinical trials.

We are only starting to understand the (patho)physiological roles played by sPrP. Further mechanistic studies are clearly required to investigate if and how sPrP indeed supports sequestration of toxic proteopathic oligomers into respective deposits, and whether its interaction with those conformers in the extracellular space may induce additional effects (such as receptor binding and cellular uptake for degradation or activation of glial responses). The relevance of Aβ-associated sPrP in brain vessels also deserves a more detailed investigation. Likewise, whether stimulated shedding –at least partially– contributes to the protective effects of certain PrP-directed antibodies in current therapeutic approaches (and clinical trials ^114^) against prion diseases remains to be investigated. Whether or not sPrP, as a soluble factor drained into body fluids, such as CSF and blood, holds potential as an easily accessible diagnostic biomarker, as a reliable reporter for treatments targeting PrP expression, or even as a read-out for any ADAM10-targeting strategies in various pathophysiological processes is currently being investigated. The sPrP-specific antibodies characterized herein lay the foundation for these and other initiated and follow-up investigations.

But is it appropriate to only discuss sPrP in the context of (neuro)protective aspects? This conclusion would probably be premature and not satisfying the actual complexity. Recent reports suggest a role of sPrP in the development and drug resistance of certain human tumors ^51,52^ while others have proposed a detrimental role in neuropathological complications caused by HIV infection ^50^. Regarding the latter, “soluble PrP” was found increased in body fluids of HIV patients with neurocognitive impairment ^125^, suggesting that our sPrP-specific antibodies could foster new systematic insight with regard to pathomechanisms and diagnostic potential beyond the field of protein misfolding diseases. It is tempting to speculate that these instances may be connected with the known harmful upregulation of ADAM10 in tumorigenesis, metastasis and inflammatory conditions ^124^.

And what about prion diseases? In murine disease models (and possibly dependent on the actual prion strain under investigation), aggregates of misfolded PK-resistant yet ADAM10-cleaved forms of PrP (sPrP^res^) were shown in recent studies using our sPrP-specific antibody for mouse samples ^41,42^. This fits earlier reports showing that misfolded PrP can, in principle, be released from cells by ADAM10 (yet, remarkably, not by phospholipases cleaving within the GPI anchor structure) ^22,126,127^. Alternatively, shed PrP could undergo misfolding in the extracellular space, similar to what was shown in transgenic mice expressing anchorless PrP ^128,129^. Notably, although ADAM10 expression in prion-infected transgenic mice appeared to correlate with reduced overall prion conversion and longer survival (indicative of reduced PrP-associated neurotoxicity) ^26,34,35^, histological assessment pointed towards ADAM10 supporting spatiotemporal spreading of neuropathological hallmarks within the brain ^35^. This dual role fits the previously described mechanistic uncoupling of prion formation/infectivity on the one hand, and neurotoxicity on the other hand (with the latter primarily being determined by cell surface PrP^C^ levels and defining disease tempo) ^130,131^. Thus, possibly depending on prion strain and the affected species, sPrP^res^ (or ‘shed prions’) may also contribute to prion spread inside and outside an organism. In this regard, it will be particularly interesting to study whether ‘proteolytic shedding’ and potential presence of sPrP^res^ in saliva, nasal secretions, urine or feces of deer and elk contributes to the efficient ‘environmental shedding’ and, hence, horizontal transmission of prions causing highly contagious CWD ^132–134^. Detailed investigations on the role of shedding in naturally occurring prion diseases in humans and other relevant mammals are certainly warranted and will profit from the findings and research tools presented herein.

## Material and Methods

### Samples and ethics statements

Postmortem human tissues (FFPE blocks and frozen samples) were acquired in the framework of diagnostic hospital and reference center activities. Use of such tissue samples (following data protection-conform anonymization) after conclusion of diagnostic procedures was in agreement with §12 of the Hamburg Hospital Act (*Hamburgisches Krankenhausgesetz*; HmbKHG) and regulations at the University Medical Center Hamburg-Eppendorf, and follows ethical regulations of the 1964 Declaration of Helsinki, its later amendments or comparable ethical standards; CSF samples: approved by the institutional review board of the independent ethics committee of the Hamburg Chamber of Physicians (PV3392).

Tissue samples (Fig. 6b-e) were obtained for diagnostic purposes from mandatory autopsies under surveillance rules and in agreement with local ethic guidelines. They were provided upon anonymization by Prof. Dr. Mara Popović (Institute of Pathology, Faculty of Medicine, University of Ljubljana, Slovenia), Prof. Dr. Herbert Budka (Division of Neuropathology and Neurochemistry, Medical University Vienna, Austria) and Prof. Dr. James Ironside, Neuropathology Laboratory, National CJD Surveillance Unit, University of Edinburgh, Edinburgh, UK.

Frozen brain samples (Fig. 7a) were obtained from the Institute of Neuropathology Brain Bank (now a branch of the HUB-ICO-IDIBELL Biobank) Hospitalet de Llobregat, Barcelona, Spain. This procurement was carried out in compliance with Spanish biomedical research regulations, including *Ley de la Investigación Biomédica* 2013 and *Real Decreto de Biobancos* 2014, and approval of the Ethics Committee of the Bellvitge University Hospital (HUB) PI-2019/108.

Human embryonic NSCs were purchased from Thermofisher, Approval number NOR/REC R0500096A (French Biomedicine Agency). The human iPSC cells used for organoids were purchased from ATCC and de-identified before being provided to the researchers at the NIH. Their use has been reviewed and determined to be exempt from IRB review; Approval number: 17-NIAID-00212, 3^rd^ of August 2017.

#### Animal samples

No experiments on living animals were performed for this study, as in accordance with the ‘3R’ principles in animal research, we profited from samples already available from previous studies. Breeding and sacrification for the sake of organ removal were approved by the ethical research committees of respective national/local authorities: *Freie und Hansestadt Hamburg – Behörde für Gesundheit und Verbraucherschutz*, Hamburg, Germany (ORG1023 for WT (C57BL/6J) and PrP-KO (*Prnp*^0/0^) mice); *Schleswig-Holsteinisches Ministerium für Energiewende, Landwirtschaft, Umwelt und ländliche Räume*, Kiel, Germany (V241-25481/2018(30-3/16) for the 5xFAD mouse). Frozen brains (used for western blot analyses) of transgenic mouse lines tg338 (sheep-VRQ), tg501 (goat-ARQ), tg110 (bovine), tg340 (human-MM129), and tg361 (human-VV129) were obtained from the breeding colony of *Istituto Superiore di Sanità*, Rome, Italy. These lines are on a *Prnp*^0/0^ background and homozygous for the transgene. Approved and supervised by the Service for Biotechnology and Animal Welfare of the *Istituto Superiore di Sanità* and authorized by the Italian Ministry of Health (decree number: 1119/2015-PR). All procedures were carried out in accordance with the Italian Legislative *Decree* 26/2014 and European Union (EU) Council directives 86/609/EEC and 2010/63/EU. Samples of prion-infected transgenic mice (Fig. 6 and Suppl. Fig. 8): Procedures were in compliance with institutional and French national guidelines and with aforementioned EU directives. Experiments were approved by the Committee on the Ethics of Animal Experiments of INRAe Toulouse/ENVT (Permit Number: 01734.01).

Brains of BSE-infected cattle and a Scrapie-infected sheep (Fig. 6c) were archived veterinarian samples of naturally occurring prion diseases.

Frozen animal brain samples (Suppl. Fig. 5) were obtained from postmortem examination in the framework of diagnostic activities of the National Institute for Agricultural and Veterinary Research (INIAV), Portugal. Use of these samples upon finished diagnostic procedures was reviewed and approved by the Quality and Environment Office of the INIAV. Samples were non-biohazardous and non-infectious.

#### Cell lines

All mammalian cell lines used in this study are listed below (Table 2). For human NSCs and iPSCs see text above.

**Table 1:** Details on human samples used in this study.

| ID | Sex | Age (y) | Diagnosis / Staging / Other | PMI | Method | Fig. | Braak stage |
| --- | --- | --- | --- | --- | --- | --- | --- |
| 1 | ♀ | 85 | control, non-ND, Cx, Cb | n.a. | IHC | 6 | n.a. |
| 2 | ♂ | 67 | sCJD (MM2C type) | n.a. | IHC | 6, S7 | n.a. |
| 3 | ♂ | 60 | sCJD (MV2K type) | n.a. | IHC | 6, S7 | n.a. |
| 4 | ♀ | 70 | sCJD (VV2 type) | n.a. | IHC | S7 | n.a. |
| 5 | ♀ | 64 | sCJD (with Kuru plaques) | n.a. | IHC, IF | 6 | n.a. |
| 6 | ♂ | 62 | vCJD | n.a. | IHC, IF | 6 | n.a. |
| 7 | ♀ | 41 | GSS (P102L mutation) | n.a. | IF | 6 | n.a. |
| TMA 1 | ♂ | 67 | control, non-ND | n.a. | IHC | 7 | - |
| TMA 2 | ♂ | 69 | AD (CERAD C), mild CAA | n.a. | IHC | 7 | IV |
| TMA 3 | ♂ | 69 | control, non-ND, brain metastasis | n.a. | IHC | 7 | - |
| TMA 4 | ♀ | 98 | AD (CERAD C) | n.a. | IHC | 7 | V |
| AD1 | ♀ | 64 | Trisomie 21 (CERAD C), front. Cx | n.a. | IF | 7 | V |
| AD2 | ♀ | 76 | AD (CERAD C), CAA, disease onset at 69 y, occip. Cx | 8 h | BVI, IF, IHC | 7, S9 | VI |
| AD3 | ♀ | 61 | AD (CERAD C), CAA, disease onset at 55 y, occip. Cx | 4 h | BVI, IF, IHC | 7, S9 | VI |
| Control A1 | ♂ | 61 | non-ND, stroke | 4 h | WB | 7 | - |
| Control A2 | ♂ | 78 | non-ND, stroke | 2 h | WB | 7 | - |
| Control A3 | ♀ | 75 | non-ND, stroke | 3 h | WB | 7 | - |
| Control A4 | ♂ | 67 | non-ND, stroke | 5 h | WB | 7 | - |
| Control A5 | ♂ | 85 | non-ND, stroke | 6 h | WB | 7 | - |
| Control A6 | ♂ | 64 | non-ND, mesial sclerosis | 3 h | WB | 7 | - |
| Control A7 | ♂ | 71 | non-ND, no neuropathol. findings | 12 h | WB | 7 | - |
| Control A8 | ♂ | 52 | non-ND, no neuropathol. findings | 5 h | WB | 7 | - |
| AD B1 | ♂ | 63 | AD | 4 h | WB | 7 | I |
| AD B2 | ♀ | 74 | AD (CERAD A) | 3 h | WB | 7 | I |
| AD B3 | ♀ | 57 | AD (CERAD A) | 5 h | WB | 7 | I |
| AD B4 | ♀ | 75 | AD | 5 h | WB | 7 | II |
| AD B5 | ♂ | 66 | AD | 21 h | WB | 7 | II |
| AD B6 | ♀ | 72 | AD | 8 h | WB | 7 | II |
| AD B7 | ♀ | 77 | AD | 3 h | WB | 7 | II |
| AD B8 | ♂ | 65 | AD | 15 h | WB | 7 | II |
| AD B9 | ♂ | 57 | AD | 4 h | WB | 7 | II |
| AD D1 | ♀ | 83 | AD | 4 h | WB | 7 | V |
| AD D2 | ♂ | 79 | AD | 5 h | WB | 7 | V |
| AD D3 | ♂ | 75 | AD | 11 h | WB | 7 | V |
| AD D4 | ♀ | 85 | AD (CERAD C) | 16 h | WB | 7 | V |
| AD D5 | ♂ | 77 | AD (CERAD C) | 8 h | WB | 7 | V |
| AD D6 | ♀ | 72 | AD (CERAD C) | 9 h | WB | 7 | V |
| CSF<sub>1</sub> | ♀ | 78 | non-ND, normal pressure hydrocephalus | biopsy | WB | 7 | - |
| CSF<sub>2</sub> | ♀ | 46 | non-ND, cerebral gliosis, migraine | biopsy | WB | 7 | - |

Abbreviations: AD = Alzheimer's disease; BVI = brain vessel isolation; CAA = cerebral amyloid angiopathy; Cx = Cortex; ND = neurodegeneration; IF = immunofluorescence analysis; PMI = postmortem interval; WB = SDS-PAGE and western blot analysis; S = Suppl. Figure

**Table 2:** List of cell lines used in this study.

| Name | Origin<br>(species, tissue) | Source | Procedures |
| --- | --- | --- | --- |
| **N2a WT** | Murine, neuroblastoma | Institute of Neuropathology, UKE, Hamburg (based on ATCC [CCL-131] ordered via DSMZ [ACC 148]) | - |
| **N2a PrP KO** | Murine, neuroblastoma | Dr. Michael William, LMU, Munich <sup>90</sup> | Transfected with WT human PrP, and GFP-tagged versions of both species' PrPs (Fig. 5b) |
| **SH-SY5Y** | Human, neuroblastoma (differentiated from a metastatic bone tumor) | Institute of Neuropathology, UKE, Hamburg (based on ATCC [CRL-2266]) | Transfected with human and murine PrP (Fig. 5c) |
| **A549** | Human, lung carcinoma | WT, ADAM10 KO<sup>a</sup> and ADAM17 KO: S. Lichtenthaler, Munich <sup>77</sup><br>WT, ADAM10 KO<sup>b</sup>: generated/customized for the Institute of Neuropathology, UKE, Hamburg, by the company *Ubigen* | Treatment with protease inhibitors and/or PMA |
| **SHEP2 (Tet2)** | Human, neuroblastoma | Institute of Biochemistry, University of Kiel, Germany; origin: NKI/AMC Amsterdam, The Netherlands (accession # CVCL_HF70; www.cellosaurus.org); | Treatment with antibodies or protease inhibitor |
| **LN235** | Human, astrocytoma | Prof. Dr. M. E. Hegi, Neurosurgery & Neuroscience Research Center, Epalinges; Centre hospitalier universitaire Vaudois, Lausanne Switzerland (accession # CVCL_3957; www.cellosaurus.org) | Treatment with antibodies or protease inhibitor |
| **U373-MG** | Human, glioblastoma | Merck/Sigma-Aldrich, #8061901 (accession # CVCL_2818; www.cellosaurus.org) | Treatment with antibodies or protease inhibitor |

**Table 3:** List of antibodies used in this study.

| Name/target | Description | Application | Company / Source |
| --- | --- | --- | --- |
| sPrP<sup>G227</sup> | Rb polycl. specific for mouse/rat sPrP | WB, IHC, IF | UKE Hamburg |
| POM1 | Ms monocl. anti-PrP (C-term. half) | T, WB | Millipore |
| POM2 | Ms monocl. anti-PrP (N-term. half) | T, WB | Millipore |
| 3F4 | Ms monocl. anti-human PrP (central) | T, WB, IHC, IF | Millipore |
| 6D11 | Ms monocl. anti-PrP (central) | T, WB | BioLegend |
| SAF84 | Ms monocl. anti-PrP / anti-PrP<sup>Sc</sup> | IHC | Cayman Chemicals |
| SAF70 | Ms monocl. anti-PrP / anti-PrP<sup>Sc</sup> | IF | Cayman Chemicals |
| β-actin | Ms monocl. anti-β-actin, clone C4 | WB | Millipore |
| β-Tubulin III | Ms monocl. anti-β-tubulin III, clone Tuj1 | IF | Covance |
| GAPDH | Rb monocl. anti-GAPDH | WB | Cell Signaling |
| BAM-10 | Ms monocl. anti-human $\beta$ amyloid | IF, IHC | Medac-Diagnostika |
| 6E10 | Ms monocl. anti-human $\beta$ amyloid | WB, IHC, IF | BioLegend |
| APP / sAPPα | Rb polycl. anti-mouse/rat $\beta$ amyloid | WB | BioLegend |
| ADAM10 | Rb monocl. anti-ADAM10 [EPR5622] | WB | Abcam |
| ADAM17 | Rb monocl. anti-ADAM17 | WB | Abcam |
| Laminin | Rb polycl. anti-human Laminin | IF | Sigma-Aldrich |
| Lectin GS-II | Alexa 488 conj. anti-human Lectin | IF | ThermoFischer Scientific |
| Iba1 | Rb polycl. anti-Iba1 | IHC | Wako Chemicals |
| GFAP | Ms monocl. anti-GFAP | IHC | DAKO |
| LAMP1 | Rat monocl. anti-Ms LAMP1 clone 1D4B | IF | Developmental Studies Hybridoma Bank (DSHB) |
| NF-L | Ms monocl. anti-NF-L | IF | Invitrogen |
| OSP | Rb polycl. anti-OSP | IF | Abcam |
| FoxG1 | Rb polycl. anti-FoxG1 | IF | Abcam |
| s100b | Rb monocl. anti-s100β | IF | Abcam |
| GABA B R1 | Ms monocl. anti GABA B R1 | IF | Abcam |
| - | Gt anti-Ms secondary / HRP | WB | Promega |
| - | Gt anti-Rb secondary / HRP | WB | Promega |
| IRDye 800 CW | Dk anti-Ms IgG | WB | Licor |
| IRDye 680 RD | Dk anti-Rb IgG | WB | Licor |
| - | Gt anti-Ms secondary / AlexaFluor-488 | IF | ThermoFischer Scientific |
| - | Gt anti-Rb secondary / AlexaFluor-647 | IF | ThermoFischer Scientific |
| - | Dk anti-Rb secondary / AlexaFluor-488 | IF | ThermoFischer Scientific |
| - | Dk anti-Rb secondary / AlexaFluor-555 | IF | ThermoFischer Scientific |
| - | Dk anti-Ms secondary / AlexaFluor-647 | IF | ThermoFischer Scientific |
| - | Ch anti-Rb secondary / AlexaFluor-647 | IF | ThermoFischer Scientific |

[WB=western blot; IHC=immunohistochemistry; IF=immunofluorescence microscopy; T=treatment; HRP= horse radish peroxidase; Ms=mouse; Rb=rabbit; Gt=goat; Dk=donkey; Ch=chicken; conj =conjugated]

### Cell culture treatments and harvesting of lysates and conditioned media

Cells were treated with 30µM of metalloprotease inhibitors GI254023X and/or GW280264X, 50nM of the ADAM10 stimulators PMA or Carbachol, or PrP-directed antibodies (as indicated) in OptiMEM for 18 h at 37°C with 5% CO_2_. Samples were then harvested for further analysis by western blot. Conditioned media was collected into cooled tubes on ice (already containing 20× pre-dissolved protease inhibitor cocktail (PI; cOmplete EDTA-free, Roche) and subsequently centrifuged at 500×*g* and 5,000×*g* for 5 min at 4°C. The remaining supernatant was used for precipitation (as described below). The confluent cells were washed with cold PBS before addition of 150 µL of RIPA buffer (50mM Tris-HCl pH 8, 150mM NaCl, 1% NP-40, 0.5% Na-Deoxycholate, 0.1% SDS) containing freshly added1×PI for 15 min on ice. Cells were scrapped, transferred to a tube and stored on ice (with short vortexing every 5 min) for 15 min. Centrifugation was performed at 12,000×*g* for 10 min at 4°C and supernatants (lysates) stored in aliquots at −80°C or mixed with water and 4× sample buffer (SB; 250mM Tris-HCl, 8% SDS, 40% glycerol, 20% β-mercaptoethanol (β-ME), 0.008% bromophenol blue (pH 6.8)), denatured at 96°C for 10 min and used for SDS-PAGE.

### Structural modeling (Peptide-protein docking) and cleavage site prediction

The 3D conformation of the human PrP peptide 217-YERESQAYYQRGS-230 was initially predicted using PEP-FOLD3 ^69^. To generate a starting template for subsequent flexible docking using the Rosetta FlexPepDock protocol ^70^, the peptide was aligned onto the crystal structure of the ADAM10 catalytic domain in complex with the C-terminal region of an adjacent ADAM10 molecule as substrate (PDB: 6BE6). Structure visualization and analysis were carried out using PyMOL (Schrödinger LLC). IceLogo plots for cleavage preferences of ADAM10 and ADAM17 were published earlier as indicated.

### Generation of sPrP-directed antibodies

The **polyclonal sPrP^Y^**^226^ **antibody** was generated (upon structural prediction of Y226 as a potential shedding site) using an anti-peptide approach following a standard 87-day polyclonal protocol (Eurogentec, Belgium). Briefly, based on the sequence information of human PrP^C^, a recombinant peptide (*N-term*-C-ESQAY**Y**-COOH; with Y-COOH representing Y226, i.e. the assumed new C-terminus of shed PrP exposed upon shedding) was produced and N-terminally coupled to *Megathura crenulata* keyhole limpet hemocyanin (KLH) as carrier protein. This immunogen was injected into two rabbits at days 0, 14, 28 and 56 of the immunization programme. Blood samples were collected from the tails at days 0, 38 and 66 for monitoring the success of the immunization process by standardized ELISA tests. Animals were sacrificed and blood collected at day 87. Standardized quality measures at Eurogentec revealed good target responses for both final bleeds and additional affinity purification was then performed on one of the sera: A second peptide (*N-term*-C-YERESQAY**Y**QRGS; representing the C-terminus of fl-PrP till the GPI anchor attachment site) was produced, coupled to a resin and served as a “negative control” to eliminate all antibodies from the polyclonal serum that would otherwise bind to fl-PrP.

Mouse **monoclonal antibody V5B2** was generated several years ago against peptide P1 (PrP214–226, CITQYERESQAYY) in BALB/c mice ^61,62^. Three groups of five BALB/c mice were injected subcutaneously on day 0 with 0.2 mg of peptide P1 bound to KLH (P1–KLH) in Freund’s complete adjuvant (0.2 mL/mouse). On days 14 and 28, the mice were injected intraperitoneally with 0.1 mg P1–KLH in Freund’s incomplete adjuvant (IFA; 0.2 mL/mouse). Blood was taken from the tail vein 10 days after the last inoculation. Antibodies against KLH, P1–KLH and peptide alone were detected in sera by indirect ELISA. A final booster dose of P1–KLH was injected on day 45 intravenously in physiological saline (0.1 mg/mouse in 0.1 mL) to mice with the highest titers of mAbs against each of the peptides. Mice were sacrificed on day 48 and their spleens removed. Splenocytes were isolated and fused with mouse NS1 myeloma cells using 50% PEG for 3 min, according to standard techniques. Cells were washed and resuspended in 96-well microtiter plates in DMEM (Dulbecco’s modification of Eagle’s medium, ICN Biomedical) supplemented with 13% bovine serum (Hy Clone) (subsequently designated DMEMbov) and with feeder cells of mouse thymocytes. The next day, DMEMbov supplemented with hypoxanthine-aminopterine-thymidine (HAT, Sigma) mixture was added to all wells. Presence of specific antibodies was determined in the supernatants after 10-14 days by indirect ELISA. Hybridomas from positive wells were transferred into larger volumes of HAT-DMEMbov and the specificity of antibodies was determined by immunohistochemistry and dot blot. Selected hybridomas were cloned in DMEMbov by the limiting dilution method and frozen in liquid nitrogen. Monoclonal antibodies were purified from the cell culture supernatants by affinity chromatography on Protein G Sepharose (Sigma), using 0.1M glycine, pH 2.7, for elution. The clone V5B2, back then, was shown to target human PrP ending at Y226 ^87^ and to detect certain disease-associated PrP forms in different prion diseases ^61,63–65^. The identity of the V5B2 target PrP226* as physiological shed PrP was revealed during the course of the study presented here.

### Antibodies

Apart from the abovementioned antibodies for the specific detection of sPrP, the following primary antibodies were employed in this study (incl. application/source):

### Recombinant prion protein production

Full-length human PrP (recPrP23-231) and truncated versions thereof (recPrP23-226, recPrP90-224, recPrP90-225, recPrP90-226, recPrP90-227, recPrP90-228, recPrP90-231) were prepared at the *Slovenian Institute for Transfusion Medicine* and expressed, purified and refolded according to our previous protocol ^89^. Plasmids encoding the variant sequences were transformed into competent *E. coli* BL21 (DE3) and grown at 37°C in 1 L of minimal medium with ampicillin (100 μg/mL), 4 g/L glucose and 1 g/L ammonium chlorid. At an OD600 of 0.8, the expression was induced with isopropyl-β-D-galactopyranoside to a final concentration of 0.8mM. Cells were harvested 12 h after induction and lysed by sonication (Q55 Sonicator, Qsonica). Inclusion bodies were washed in buffer containing 25mM Tris-HCl, 5mM EDTA and 0.8% Triton X-100 (pH 8) and then in bi-distilled water several times. The isolated inclusion bodies were solubilized in 6M GndHCl and purified on a 5mL FF Crude HisTrap column (GE Healthcare), equilibrated in binding buffer (2M GndHCl, 500mM NaCl, 20mM Tris-HCl and 20mM imidazole (pH 8)). Proteins were eluted with 500mM imidazole and dialysed against 6M GndHCl using Amicon centrifugal filters (MW cut-off: 3000 Da, Millipore). Purified proteins were stored at −80°C or refolded by dialysis against refolding buffer (20mM sodium acetate and 0.005% NaN_3_ (pH 4.5)) using SnakeSkin™ Dialysis Tubing (MW cut-off: 3500 Da, Thermo Scientific). Purified proteins were analyzed by SDS-PAGE under reducing conditions.

### ELISA

Microtiter plates (CORNING 9018, Costar) were coated with either 50 µL of recombinant human PrP C-terminally ending at Y226 (recPrP23-226; 0.5 µg/mL) or peptide ‘P1’ (5 µg/mL) in ELISA coating buffer (carbonate/bicarbonate buffer, pH 9.6), incubated overnight at 4°C, washed with PBS/ Tween20 (buffer B) and then blocked with 1% bovine serum albumin in PBS/Tween20 (buffer C) (Sigma-Aldrich). 50 µL of mouse monoclonal V5B2 or rabbit polyclonal sPrP^Y226^, titrated in buffer C, were added to the wells and incubated for 90 min at 37°C. Plates were washed in buffer B, and 50 µL of HRP-conjugated anti-mouse or anti-rabbit immunoglobulin (Jackson ImmunoResearch), both diluted 1:2000 in buffer C, were added and incubated for 90 min at 37°C. After final washes, 2,2’-azino-bis (3-ethylbenzthiazoline-6-sulfonic acid) (ABTS, Sigma-Aldrich) substrate was added to each well. Absorbance was measured at 405 nm after 10 min of incubation at 37°C in a Tecan Sunrise microplate spectrometer (Tecan).

### Human neural stem cells culture, neuronal differentiation, and treatment

We used HuPrP-overexpressing H9NSC cells (derived from human embryonic stem cells (WAO9, Wicells)). These cells were obtained following transduction of H9NSC with the pWPXL-PrP-IRES-GFP lentivirus coding for wild-type human PrP as well as GFP. These lentiviral vectors were derived from the HIV-based Tronolab vectors and produced by the Biocampus PVM Vectorology Platform. For the treatment of neurons derived from HuPrPH9NSC, amplified HuPrPH9NSC were seeded at a density of 9.6×10^5^ cells per well on 6-well plates coated with Geltrex in StemProNSC medium and placed in a 37°C, 5% CO_2_ and 5% O_2_ incubator. Three replicas for the four experimental groups were prepared. The day after seeding, the medium was replaced by a N2 + bFGF 20 ng/mL medium (KO/DMEM/F12 supplemented with 1% (v/v) N2 supplement, 1mM glutamine and 1% penicillin/streptomycin). The medium was changed every two days. bFGF was added every day to commit NSC into neuronal progenitor cells. At day 5 of differentiation, bFGF was removed and the cells were maintained for seven more days in N2 medium alone (again changed every two days). The cells were then cultivated in N2 medium containing laminin (1 ng/mL) and BDNF (10 ng/mL) until day 30 of differentiation. During this period half of the media volume was refreshed every 3 days. Total media volume per well was 1 mL. The cells were treated (condition A: control medium containing DMSO (1/5,000); condition B: ADAM10 inhibition with GI254023X (6µM); condition C: HuPrP-directed 3F4 antibody [6 µg/well]; condition D: PrP-directed 6D11 antibody [6 µg/well]) at day 29 of differentiation and incubated for 18 h. Conditioned media and cells were then collected as follows: concentrated PI cocktail dissolved in 30 µL PBS was first filled in the collection tube (low-binding; Eppendorf) and conditioned media was carefully aspirated from the cell layer, added to the tube on ice and mixed by gentle inverting. Two mild centifugations of 5 min at 4°C (500×g; 5,000×g) were performed to remove cell debris. For cell lysis we freshly dissolved one tablet each of PI and PhosStop (Roche) in 8 mL of RIPA-Buffer. After carefully washing the cells on ice two times with cold PBS, 120 µL of this RIPA-Buffer were added, lysis was mechanically supported by scratching off the cells from the dish and pipetting up and down. Total duration of lysis was 20 min. Each sample (media and lysates) was stored at −80°C until analysis.

### Human iPSC-derived cerebral organoids

#### Organoid generation and culture

Human induced pluripotent stem cell (iPSC) line KYOU-DXR0109B (ATCC) was routinely cultured on growth-factor reduced Matrigel (Roche) in mTeSR1 culture medium (StemCell Technologies) under standard incubator conditions (humidified, 37°C, 5% CO_2_). Cerebral organoids were generated from iPSCs using the STEMdiff cerebral organoid kit (StemCell Technologies) as per the manufacturer’s instructions. For long term culture they were maintained in cerebral organoid media (1× glutamax, 1× penicillin-streptomycin solution, 0.5× non-essential amino acids, 0.5% (v/v) N2, 1 μL/4 mL insulin, 1% (v/v) B12 plus retinoic acid and 1 μL/286 mL β-ME in 1:1 Neurobasal:DMEM-F12 medium) on an orbital shaker at 85 rpm in a standard tissue culture incubator. This procedure was based on the protocol by ^135^.

#### Organoid imaging

Brightfield images of overall organoid morphology were captured using a Leica DMIL LED inverted microscope with a Leica HC 170 HD digital camera. Moreover, cerebral organoids were prepared for frozen sectioning by fixation in 10% (w/v) formalin for 24 h at room temperature (RT). Following washing in PBS, fixed organoids were incubated in 20-30% (w/v) sucrose for 24 h at RT, and then frozen at −20°C in optimal cutting temperature medium (Ted Pella Inc). For immunofluorescence stainings, slices were permeabilised in 0.1% (v/v) Triton-X-100 for 10 min, then blocked in 10% (v/v) FBS, 0.1% (w/v) BSA in PBS for 30 min before staining with primary antibodies in antibody buffer (1% (v/v) FBS, 0.1% (w/v) BSA in PBS) at the following dilutions: NF-L (Invitrogen) 1:50, OSP (Abcam) 1:50, FoxG1 (Abcam) 1:50, PrP SAF70 (Cayman Chemicals) 1:100, s100b (Abcam) 1:50. Secondary antibodies anti-rabbit-AlexaFluor-488 and anti-mouse-AlexaFluor-647 were diluted 1:250 in antibody buffer. Slides were mounted in Fluoromount-G Mounting Medium containing nuclear stain DAPI (Invitrogen). Images were captured using an EVOS FL Auto (Invitrogen) widefield fluorescence microscope system.

#### Organoid treatments and sample harvesting

For experimental treatments, organoids were transferred into 24-well low adhesion plates (Corning) in 1 mL phenol red-free OptiMEM (Gibco) with reduced cerebral organoid media supplements (10% of routine culture concentrations) and the plates were incubated on the orbital shaker as for routine culture. For the GI254023X treatments (10µM), organoids were pre-incubated with the compound for 2 h before changing media to fresh OptiMEM (already containing the compound) for 24 h. Anti-PrP 3F4 antibody and anti-mouse secondary antibody control (8 µg per well) treatments were set up in 100 µL of OptiMEM for one hour before diluting into the final media volume (1 mL) for 24 h.

Culture media collected from the treatments was supplemented with a 10× concentrated solution of PI cocktail (Roche) at a 9:1 conditioned:fresh media ratio (i.e., 1× final concentration of PI), then centrifuged at 500×g for 10 min at 4°C to remove residual cell debris. Organoid lysates were made based on the wet weight of each organoid. Sufficient RIPA lysis buffer (Pierce) with 1× PI was added to make a final 10% (w/v) homogenate and organoids were triturated. Conditioned media and lysates were stored at <-20°C until further assessment.

### Biochemical methods

#### Trichloroacetic acid (TCA) precipitation

For the precipitation of proteins from cell culture supernatants, serum-free (OptiMEM) conditioned media of the overnight cultures were used. 10 µL (1/100 of vol.) of 2% sodium deoxycholate (NaDOC) was added to 1 mL sample and was shortly vortexed. After 30 min of incubation on ice, 100 µL (1/10 of vol.) of TCA (6.1N, Sigma) were added to each sample, vortexed and incubated again for 30 min on ice. Samples were then centrifuged at 15,000×g for 15 min at 4°C. Next, the supernatant was entirely aspirated, and the pellet air-dried for 5 min and finally dissolved in 100 µL of 1× SB containing β-ME. Due to remaining TCA and low pH, the blue SB turns yellow; for neutralization (and recovery of the blue color) 1.5 µL of 2M NaOH were added and samples then boiled for 10 min at 96°C.

#### Immunoprecipitation (IP)

Immunoprecipitation was carried out using dynabeads (Pierce Protein A/G Beads). Briefly, media from UW476 cells, cultured in OptiMEM for 48 h, was collected. After PI addition, conditioned media was centrifuged first at 500×g and then at 5000×g (each for 10 minutes at 4°C). The resulting supernatant was transferred to new tubes. Next, 750 µL aliquots of this supernatant were divided into different tubes, each receiving 7.5 µg of different antibodies (monoclonal V5B2 and polyclonal sPrP^Y226^ for sPrP; monoclonal POM2 for total fl-PrP) or PBS (negative control). TCA-precipitated conditioned media was also added as a control for validating total sPrP amount. 40 μL of beads were washed with 200 μL of IP Lysis/Wash Buffer (provided in the kit). The antigen/antibody mixture was added to the beads and incubated for 2 h at RT in a rotator. Beads were magnetically immobilized to the tube wall and the supernatant (containing unbound proteins) was saved for analysis. Beads were washed twice with IP Lysis/Wash Buffer, followed by a wash with ultra-pure water. Samples were then eluted with Elution buffer (kit content). To neutralize the low pH, Neutralization Buffer (kit content) was added. For western blot analysis, 33 μL of 4× SB (with 5% β-ME) was added to each sample, which was then boiled for 10 min at 96°C.

#### Homogenization of human and animal tissue samples

Frozen brain tissues from human and animals were used to prepare 10% (w/v) homogenates in RIPA buffer containing PI and PhosStop (Roche). Briefly, samples were homogenized either manually with 30 strokes in a dounce-homogenizer or using in-tube beads (Precellys) and incubated on ice for 15 min, prior to centrifugation at 12,000×g at 4 °C for 10 min. Total protein content was assessed by Bradford assay (BioRad). Homogenates were either further processed for SDS-PAGE (i.e., 30 µL of 10% homogenate + 120 µL of H_2_O + 50 µL 4× SB (containing 5% β-ME); denaturation at 96°C for 10 min) or stored at −80°C. CSF samples were stored at −80°C, gently thawed on ice, mixed 1:3 with 4× SB (containing 5% β-ME) and boiled for 10 min at 96°C.

#### SDS-PAGE and western blotting

15-30 μL of denatured samples in SB (tissue homogenates, cell lysates, or precipitated conditioned medium) were loaded on precast Nu-PAGE 4 to 12% Bis-Tris protein gels (Thermo Fisher Scientific). After electrophoretic separation, wet blotting (at 200 mA per gel for 1 hour) was performed to transfer proteins onto 0.2 µm nitrocellulose membranes (Bio-Rad). Total protein staining was performed according to the manufacturer’s protocol (Revert™ Total Protein Stains kit; Licor). Thereafter, membranes were blocked for 45 min with either 1× RotiBlock (Carl Roth) in Tris-buffered saline containing 1% Tween 20 (TBS-T) or 5% skimmed dry milk (dissolved in TBS-T) under gentle agitation at RT. Membranes were incubated overnight with the respective primary antibodies in the corresponding blocking reagents at 4°C with gentle agitation. The following day, membranes were washed four times with TBS-T and incubated for 1 h at RT with either HRP-or IRDye-conjugated secondary antibodies (Licor). After several washes with TBS-T, membranes were developed [after incubating blots for 5 min with either Pierce ECL Pico or SuperSignal West Femto substrate (Thermo Fisher Scientific)] with a ChemiDoc imaging station (Bio-Rad) or were scanned using the Odyssey DLx imaging system (Licor). Densitometric quantification was done using the Quantity One software (Bio-Rad) and Image studio lite version 5.2 (Licor) followed by further analysis in Microsoft Excel and GraphPad Prism software.

### (Immuno)histochemical stainings and immunofluorescence analyses

#### Immunohistochemistry (IHC)

Formaline-fixed paraffin-embedded (FFPE) brain tissues were used for immunohistochemical stainings. Samples from patients or animals with prion disease were incubated in formic acid (98-100%; duration depending on samples size) prior to embedding. Sections of 4 μm were prepared with a microtome and submitted to immunostaining following standard IHC procedures using a Ventana BenchMark XT machine (Roche Diagnostics). Sections were deparaffinated and underwent antigen retrieval by boiling for 60 min in 10mM citrate buffer (pH 6.0). Afterwards, sections were incubated with primary antibodies diluted in 5% goat serum (Dianova, Hamburg, Germany), 45% TBS (pH 7.6), 0.1% Triton X-100 in antibody diluent solution (Zytomed, Berlin, Germany) for 1 h. Primary antibody (for further information refer to list above) dilutions were: V5B2 (1:50) or sPrP^Y226^ (1:50), SAF84 (1:100, for PrP^C^ and PrP^Sc^; note that for the latter (Fig. 6a) a harsh pretreatment with formic acid (5 min) followed by 30 min at 95°C in 1.1mM sodium citrate buffer [2.1mM Tris-HCl and 1.3mM EDTA (pH 7.8)], 16 min in PK and 10 min in Superblock was performed). Secondary antibody treatment was performed using anti-rabbit or anti-mouse Histofine Simple Stain MAX PO Universal immunoperoxidase polymer or Mouse Stain Kit (for detection of mouse antibodies on mouse sections). Detection of antibodies was done by Ultra View Universal DAB Detection Kit (brownish signals) or Ultra View Universal Alkaline Phosphatase Red Detection Kit (yielding pink signals) using standard machine settings (all solutions were from Ventana, Roche). Counterstaining (light blue background) was done according to standard procedures.

Stained sections were inspected, and representative pictures taken in TIF format on a digital microimaging device (DMD108, Leica) or with a Hamamatsu Slide Scanner and NDP.view2 software. The final picture processing for better presentation consisted of cropping, white balancing (graduation curves; for IHC) and brightness adjustment (equally to all channels; for IF; see below) performed with Adobe Photoshop Elements 15 during figure assembly without affecting the findings and conclusions.

#### Immunofluorescence (IF) stainings of FFPE sections

Paraffin tissue sections were cut at 3 μm and thoroughly deparaffinized (2× 20 min in Xylol and a descending alcohol row). Antigen retrieval was performed by boiling the sections in Universal R buffer (#AP0530-500; Aptum) for 20 min. Sections were briefly rinsed and blocked for 1 h. Antibodies against sPrP (1:100) and amyloid β (Aβ; 1:100; BAM-10 (Fig. 7c human sample) were incubated overnight at 4°C. After intensive washing, AlexaFluor488- and AlexaFluor555-coupled anti-rabbit and anti-mouse secondary antibodies were applied for 1.5 h. Sections were washed again and mounted in DAPI-Fluoromount-G (SouthernBiotech). Data acquisition was performed using a Leica Sp5 confocal microscope and Leica application suite software (LAS-AF-lite).

The V5B2 epitope distribution in prion plaques (Fig. 6d,e) was studied using indirect IF on 5 μm-thick sections of FFPE cerebellar samples from patients with different prion diseases. Briefly, freshly deparaffinated sections were subjected to antigen retrieval, involving 30 min autoclaving at 121°C in distilled water and 5 min incubation in 96% formic acid. After rinsing, 4% normal horse serum in buffer (block) was applied (20 minutes), followed by incubation with either V5B2 or 3F4 monoclonal antibody (at 20 μg/ml, overnight, at RT). Biotinylated horse anti-mouse secondary antibody (1:1,000, 90 min, Vector Laboratories) was applied, followed by incubation with streptavidin-Alexa 488 (1:750, 90 min, Molecular Probes). Next, fluorescence microscopy of single-labeled samples and image collection was performed, followed by second labeling: 4% normal donkey serum block was followed by V5B2 or 3F4 monoclonal antibody incubation (20 μg/ml, overnight, at RT). Signal detection was performed using Alexa 546-conjugated donkey anti-mouse secondary antibody (1:1,000, 90 min, Molecular Probes). A Nikon Eclipse E600 fluorescent microscope equipped with appropriate filters (EX 465-695, DM 505, BA 515-555 and EX 528-553, DM 565, BA 590-650) and a Nikon DXM 1200 digital camera was used for fluorescence microscopy. Alternatively, Leica TCS confocal microscope (SP2 AOBS; Leica Microsystems) was used employing the 488 nm line of the Argon laser and 543 nm Helium-Neon laser excitation light through an acousto-optical beam splitter (Leica Microsystems). The emitted light was detected at 500-540 nm (green) and 543 nm (red) using spectrophotometer (SP2, Leica Microsystems). The excitation crosstalk was minimized by the sequential scanning. Images were processed using Leica confocal software.

#### IF staining of free-floating sections

Brains (Fig. 7c murine sample) were postfixed for another 4 h in 4% PFA (in PB) and then incubated in 30% (w/v) sucrose solution (in PB). After sinking down, brains were cut with a Leica 9000 s sliding microtome (Leica) into 35 µm thick free-floating sections. For IF staining, sections were incubated in blocking solution (0.5% Triton-X 100, 4% normal goat serum in 0.1M PB pH 7.4) for 2 h at RT, followed by incubation with the primary antibody/antibodies (6E10 for human Aβ, LAMP1, sPrP^G227^) in blocking solution at 4°C overnight. Sections were washed three times with washing solution (0.1M PB pH 7.4, 0.25% Triton X-100), incubated for 90 min in secondary antibody (in washing solution), washed two times again in the washing solution and one time in washing solution without Triton X-100. Finally, sections were mounted on glass slides, embedded in Mowiol (DABCO) and analyzed with a Zeiss LSM 980 fluorescence microscope equipped with an automated stage and the ZEN 3.3 software (Zeiss).

#### Brain vessel isolation and respective IF staining

Human brain microvessels were isolated as previously described ^136^. The tissue was homogenized in 1 mL MCDB131 medium (ThermoFischer Scientific) using a dounce homogenizer, further diluted in medium and centrifuged (4°C) at 2,000×g for 5 min. The pellet was resuspended in 15% (w/v) 70 kDa dextran and centrifuged (4°C) for 15 min at 10,000×g. The microvessel-containing pellet was retrieved and transferred to a 40 µm cell strainer. Isolated microvessels were fixed on the cell strainer with 4% PFA/PBS, retrieved in 1% BSA/PBS and centrifuged (4°C) for 10 min at 2,000×g. The pellet was dissolved in PBS and applied to Superfrost microscope slides. After air drying, the slides were stored at −80°C. Isolated 2D microvessels were stained with lectin GS-II (1:200), sPrP^Y226^ (1:500) and 6E10 (1:200) or laminin (1:30),V5B2 and Thioflavin (1:200). High-resolution images were obtained with a Leica TCS SP8 confocal laser scanning microscope (Leica Microsystems) using a 63× immersion oil lens objective.

## Acknowledgements

We thank Kristin Hartmann, Edda Thies, and Celina Soltwedel as well as the Mouse Pathology Core and the Microscopy Imaging Facility ‘UMIF’ (all at UKE Hamburg, Germany), and Bradley Groveman (NIH Hamilton, USA) for excellent technical assistance. N2a cells depleted for PrP (PrP-KO) were kindly provided by Dr. Michael Willem (LMU, Munich, Germany) and LN235 cells by Prof. Dr. Monika E. Hegi (University of Lausanne, Centre hospitalier universitaire vaudois, Epalinges/Lausanne, Switzerland). Some human brain tissue samples were kindly provided to BTCS (upon completion of diagnostic procedures and full anonymization) by Prof. Dr. Mara Popović (Institute of Pathology, University of Ljubljana, Slovenia), Prof. Dr. Herbert Budka (Medical University Vienna, Austria) and Prof. Dr. James Ironside (National CJD Surveillance Unit, University of Edinburgh, UK). We also thank R-Biopharm (Darmstadt, Germany) for providing V5B2-stained human, sheep and cattle brain images (in the framework of a former Collaboration Agreement with BTCS).

