## Supplemental Figures 1-9 for "Cleavage site-directed antibodies reveal the prion protein in humans is shed by ADAM10 at Y226 and associates with misfolded protein deposits in neurodegenerative diseases"

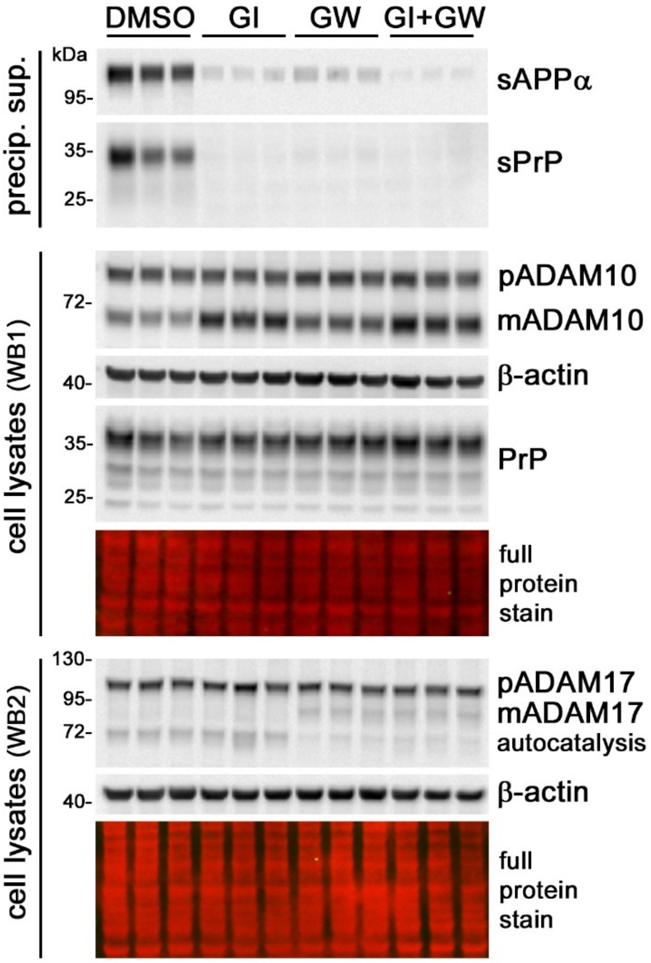


**Suppl. Fig. 1:** Western blot (WB) analysis of sPrP and sAPPα (in TCA-precipitated conditioned media) and PrP, premature (p) and mature/active (m) ADAM10 (WB1) and ADAM17 (WB2) in lysates of the human glioblastoma-derived cell line U373-MG. Cells were treated with metalloprotease inhibitors GI254023X (GI) or/and GW280264X (GW) or with the diluent only (DMSO; as control). β-actin and total protein staining served as loading controls. Note that, as in A549 cells (Fig. 1c), GI alone does not inhibit ADAM17 activity (as judged by the lack of inhibition of a previously reported postlysis autocatalytic processing step [Ref. [79]]), whereas its inhibitory effect on ADAM10 is sufficient to abolish PrP shedding

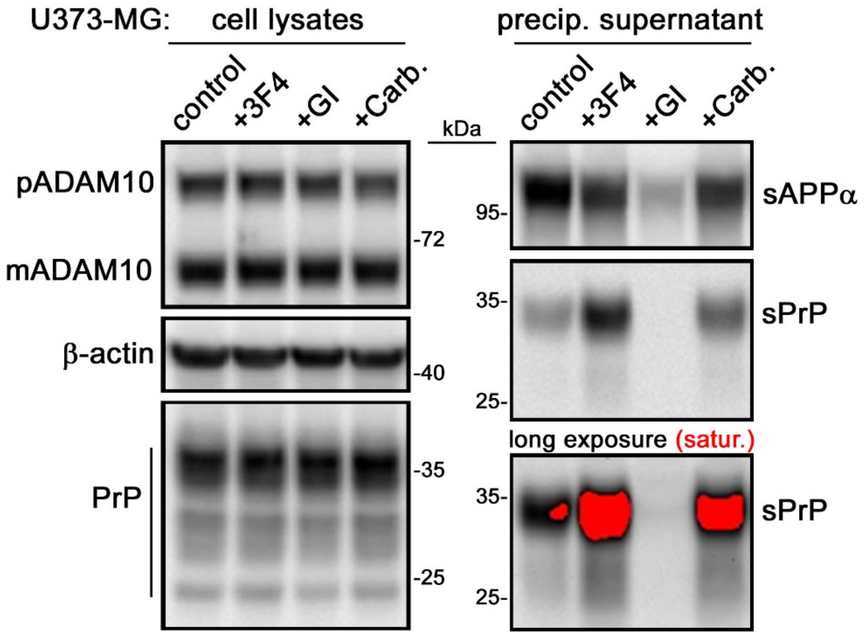


**Suppl. Fig. 2:** Western blot analysis of human U373-MG cell lysates (blots on the left) and respective precipitated conditioned media supernatants (on the right). Cells were treated o.N. with the ADAM10 inhibitor GI or with either PrP-directed IgG (+3F4) or the compound Carbachol (+Carb.) to stimulate PrP shedding. While GI treatment only reduced sAPPα levels (likely due to residual ADAM17 activity compensating as alternative APP α-secretase for inhibition of ADAM10), it completely abolishes PrP shedding. Red signal indicates saturation upon long exposure densitometric detection of the sPrP blot

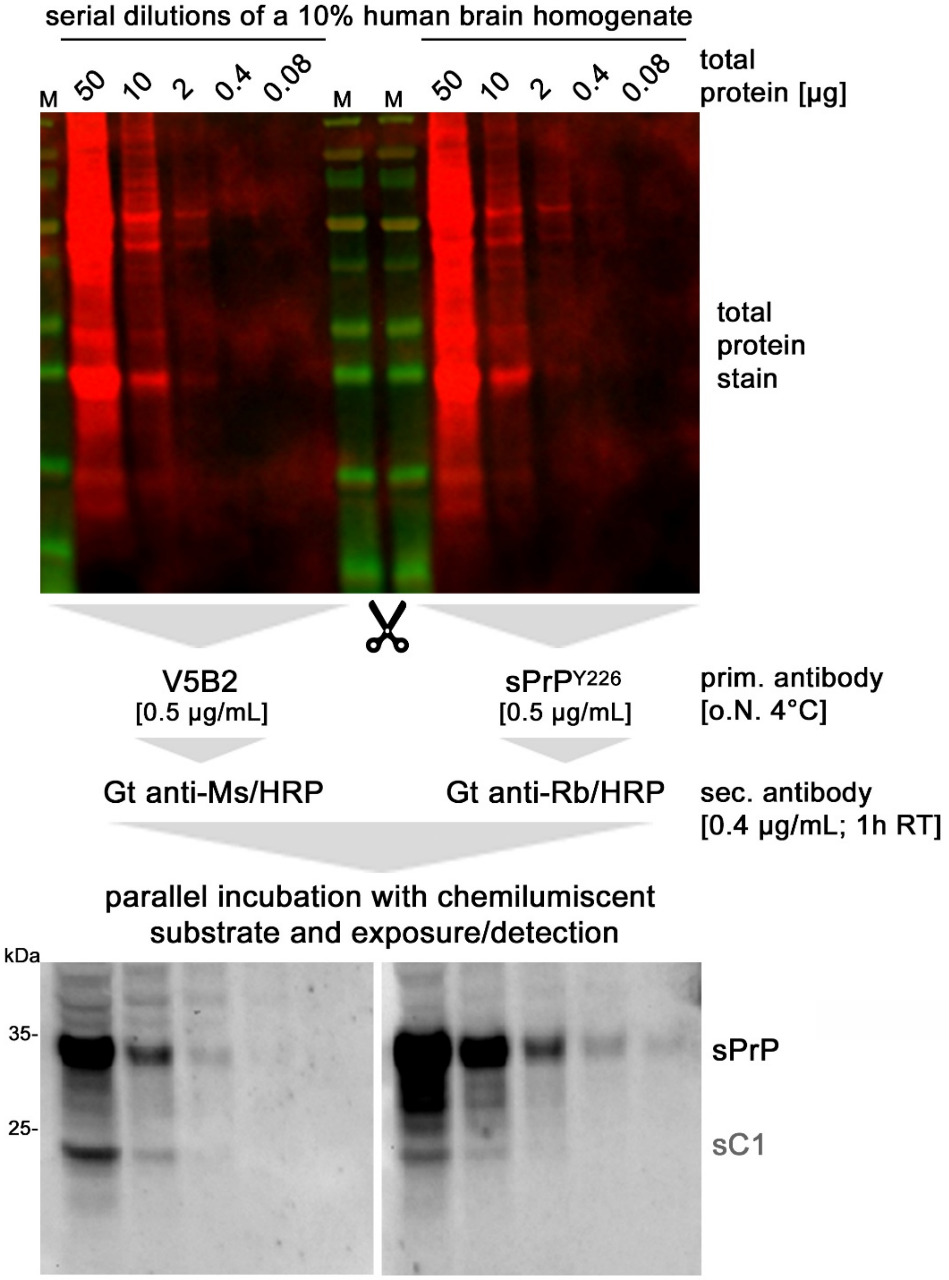


**Suppl. Fig. 3:** Immunoblot analysis directly comparing monoclonal V5B2 and polyclonal sPrP$^{Y226}$ antibodies with regard to detection sensitivity towards denatured sPrP in serial dilutions of human brain homogenates. For ideal comparison, both blot parts derive from the same SDSgel and blotted membrane. After staining of total protein, the blot was cut into two parts (as indicated by the scissors symbol) for the sake of incubation with respective first and secondary (goat, Gt) antibodies (equal incubation times, equal antibody concentration, equal washing steps [as indicated]). After washing, both blot parts were re-united for incubation with chemilumiscent substrate and for parallel detection. Shed PrP and the shed C1 fragment (sC1; resulting from shedding of already α-cleaved PrP) are detected by both antibodies with differing sensitivity. M = MW marker

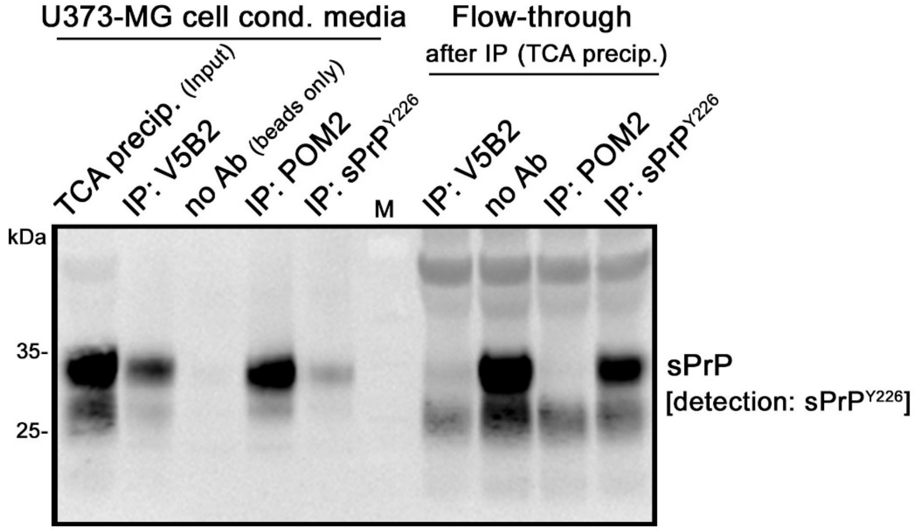


**Suppl. Fig. 4:** Immunoprecipitation (IP) of released/shed PrP from conditioned media of human U373-MG cells. Efficiency of different antibodies for pull-down of shed PrP was in the (qualitative) rank order POM2 > V5B2 > sPrP$^{Y226}$. It should be noted that POM2 has four epitopes within PrP`s disordered N-terminal domain, which may support binding of two sPrP molecules per IgG. Moreover, POM2 could pull-down full-length PrP located on extracellular vesicles (and hence sPrP molecules possibly bound to the latter). TCA precipitated media (input) and flow-through (after IP; i.e. non-bound molecules) are shown for comparison. No unspecific binding was observed and, hence, no pull-down was achieved with beads only (“no Ab“). Detection of the blot was done with the polyclonal sPrP$^{Y226}$ antibody

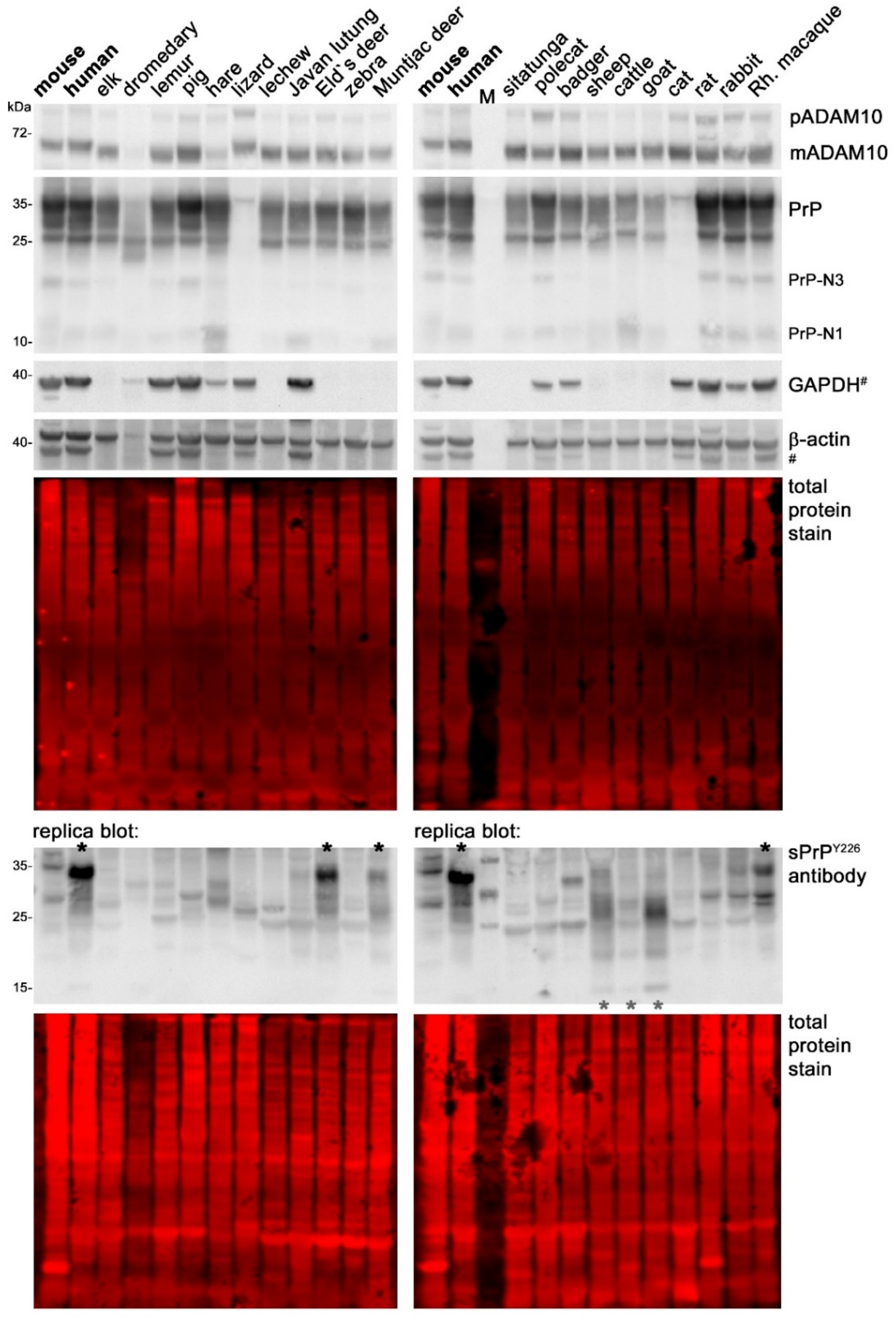


**Suppl. Fig. 5**: Immunoblot analysis of CNS tissue samples of different animals. ADAM10, total PrP (including shorter N-terminal α- and γ-cleavage fragments N1 and N3, respectively), GAPDH and β-actin were detected on the upper blots, while sPrP was detected on a replica blot. Due to sequence/epitope differences, not all proteins are detected equally in all species. This especially was the case for GAPDH, which ade us re-probing the blot with another housekeeping marker, β-actin (# indicates the previous GAPDH signals). The dromedary brain sample (4th lane) was pretty degraded at the time of assessment, and cat PrP could not be detected with the POM2 antibody used here (4th last lane). Importantly, while imperfect preservation and partial degradation of samples may be an issue here, and although unspecific bands appear upon detection with the polyclonal sPrP$^{Y226}$ antibody, a pattern similar to human sPrP appeared in the samples from Rhesus macaque as well as Eld`s and Muntjac deer (highlighted by black asterisks at the top of the blot); and a pattern reminiscent of a shed C1 fragment (see Suppl. Fig. 3) was observed in sheep, cattle and goat brain (grey asterisks at the bottom of the blot). M = MW marker

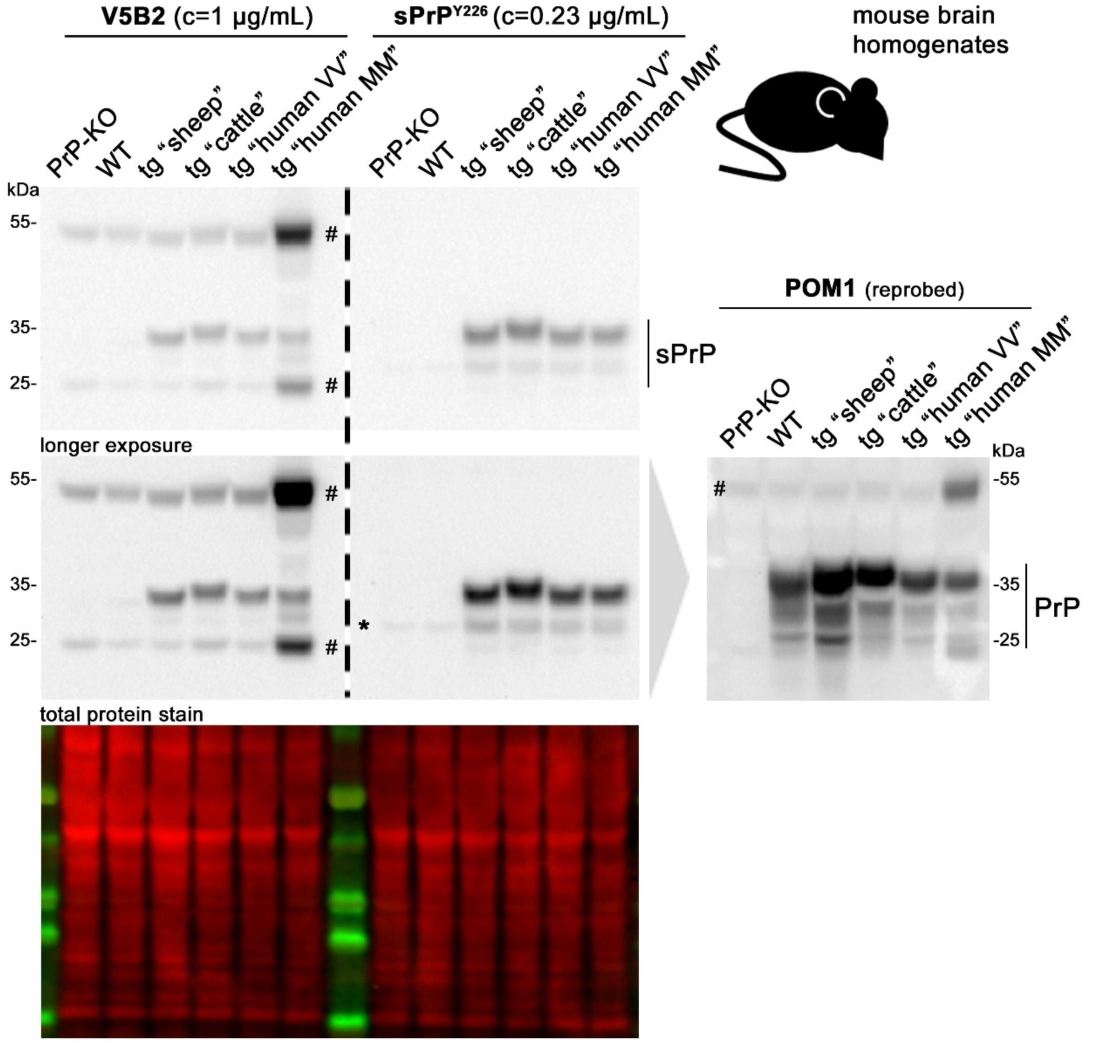


**Suppl. Fig. 6**: Comparison of monoclonal V5B2 and polyclonal sPrP$^{Y226}$ in immunoblot analysis of mouse brains. Duplicate samples were run in one gel and blotted on one membrane (see total protein stain) and only seperated (dashed line) for incubation with indicated primary (and respective secondary) antibodies. After washing, both blot parts were handled in parallel for chemilumiscent substrate incubation and detection. Both antibodies detected shed PrP only in transgenic (tg) mice expressing sheep, cattle or human PrP (for the latter, two different lines with MM or VV polymorphism at PrP position 129 were used). No specific signals were detected in PrP-KO or WT mice. Despite very similar overall results, polyclonal sPrP$^{Y226}$ revealed stronger specific signals albeit a lower antibody concentration. While detection with mouse antibodies V5B2 and (to a lesser extent) POM1 (used for re-probing/detection of total PrP) and their respective anti-mouse secondary antibodies revealed bands for IgG heavy and light chains present in the samples (indicated by #), sPrP$^{Y226}$ showed a weak PrP-independent unspecific band in PrP-KO and WT brain (running at the height of monoglycosylated sPrP in tg mice; marked by an asterisk)

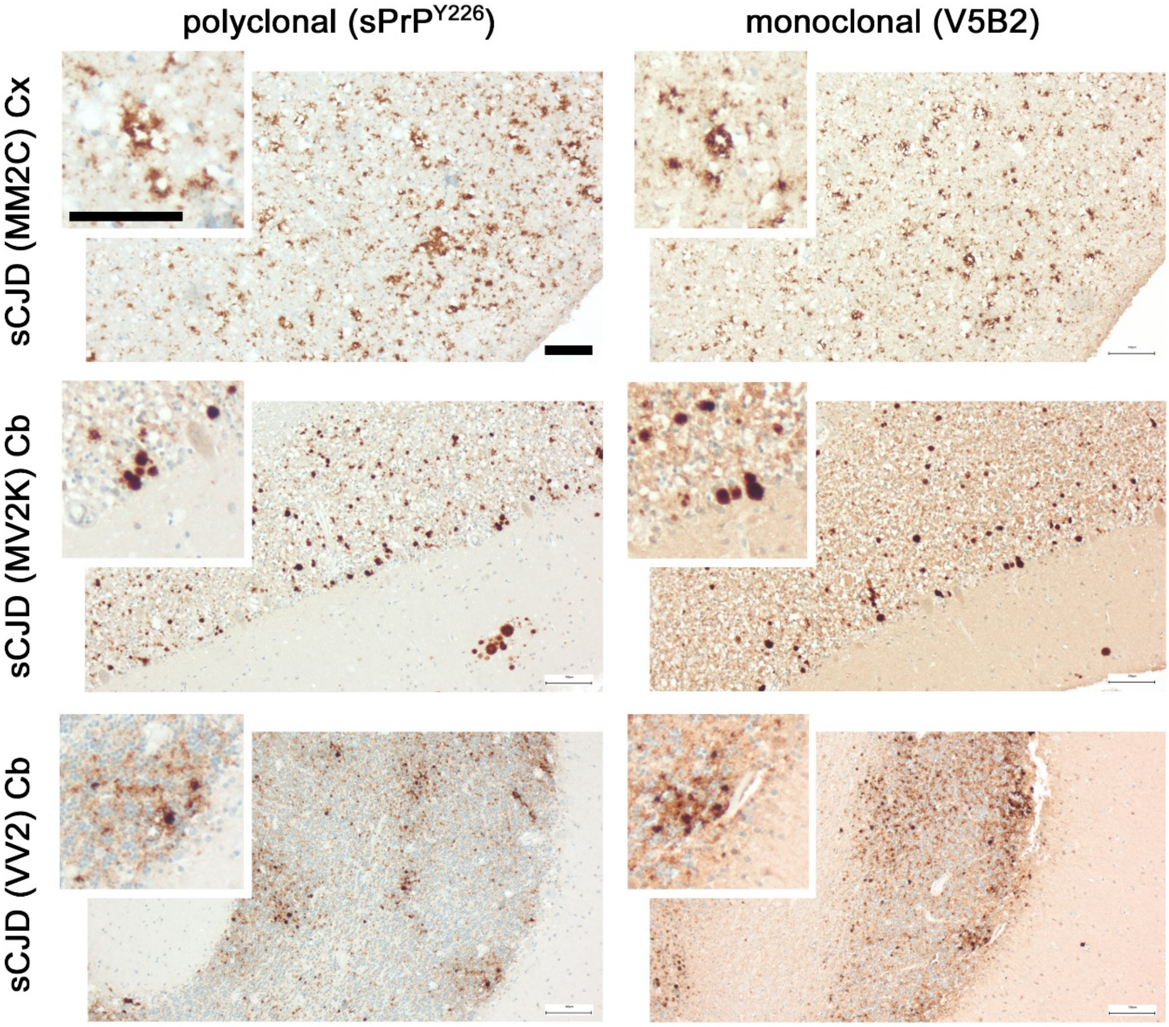


**Suppl. Fig. 7**: Comparison of polyclonal sPrP$^{Y226}$ and monoclonal V5B2 antibody in immunohistochemical assessment of brain sections of three different cases of sporadic CJD (subtype classification as indicated on the left). No PK digestion has been performed here. Cx = cortex, Cb = cerebellum. Scale bars: 100 µm

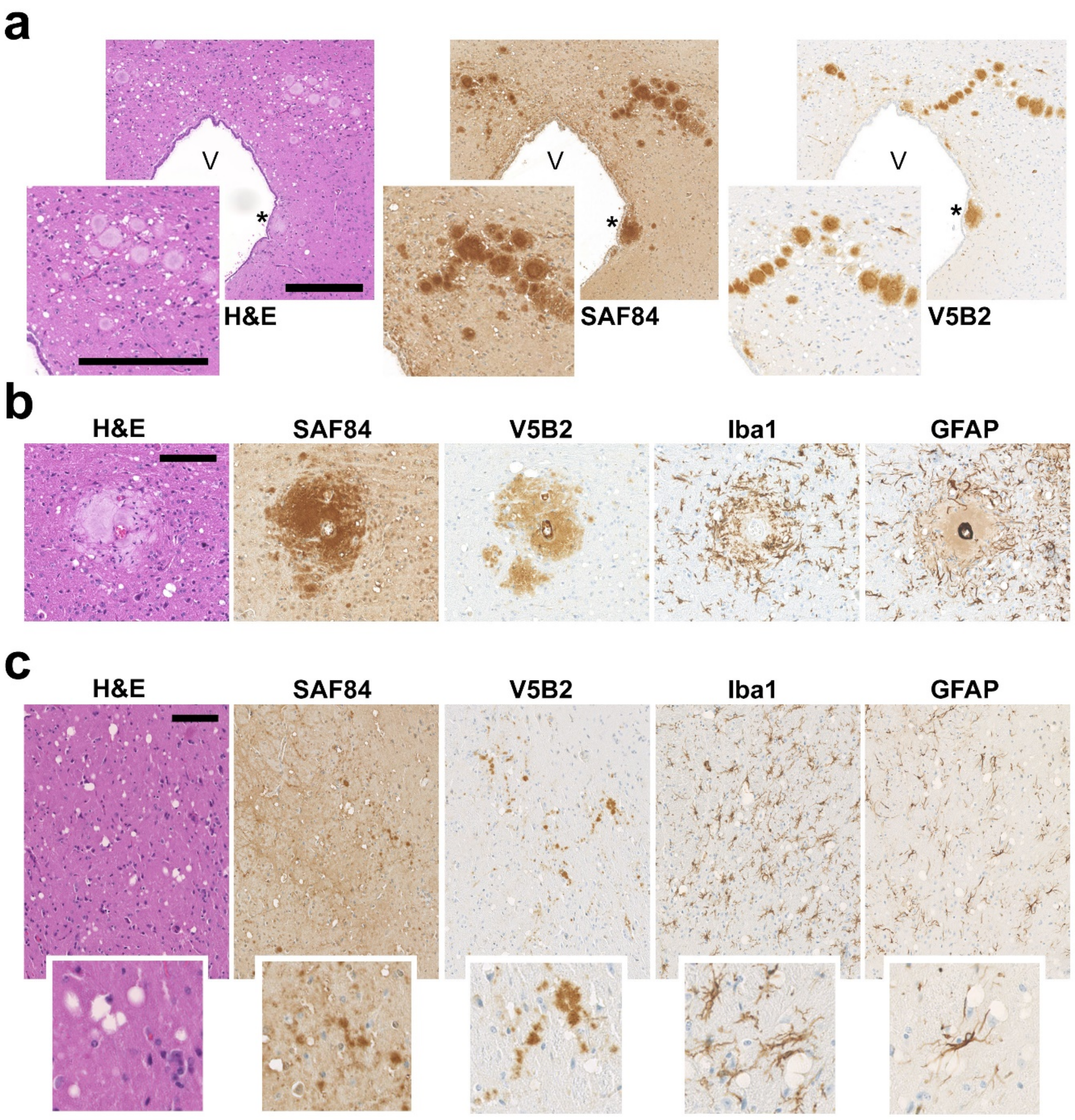


**Suppl. Fig. 8**: Immunohistochemical analyses of prion-infected transgenic mice. (**a**,**b**) Tg338 mice (expressing ovine PrP) infected with NPU1 prions show extented clusters of large prion deposits in the brain stem. Shed PrP (detected with V5B2 antibody) associates with many, yet not all of these deposits. Asterisk highlights a sPrP-positive deposit close to the ventricle (V) wall. (**b**) A representative large and amyloid-like prion deposit around a brain vessel (in the center) is positive for sPrP and surrounded by activated glia (Iba1: microglia; GFAP: astrocytes). (**c**) Prion deposits, distribution of sPrP and activated glia cells in a subthalamic area of vCJD-infected transgenic mice expressing bovine PrP. H&E staining in **a**-**c** also reveals spongiform changes. Scale bars: 250 µm (in a), 100 µm (in b,c)

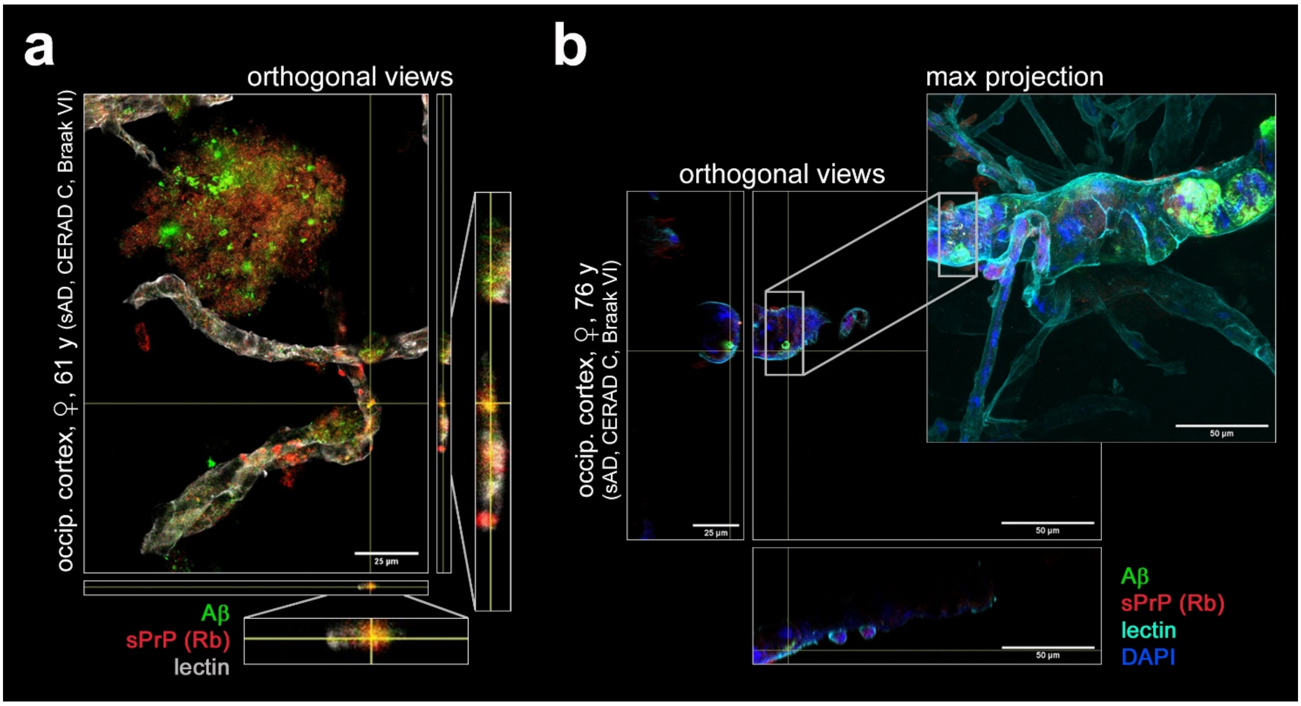


**Suppl. Fig. 9**: Confocal immunofluorescence microscopy of brain vessels isolated from human AD brain. (**a**) The same sample/analysis as in Fig. 7d, yet provided as orthogonal view representation highlighting the colocalization of sPrP (detected by polyclonal sPrP$^{Y226}$; Rb) and amyloid (Aβ) in/at brain vessels. (**b**) Orthogonal views and max projection of sPrP and Aβ in purified brain vessel of another AD patient (same as in Fig. 7e). DAPI was used to stain nuclei, lectin as an endothelial marker. Scale bars as indicated
